# Global analyses of the dynamics of mammalian microRNA metabolism

**DOI:** 10.1101/607150

**Authors:** Elena R. Kingston, David P. Bartel

**Affiliations:** Howard Hughes Medical Institute and Whitehead Institute for Biomedical Research, Cambridge, MA 02142, USA; Department of Biology, Massachusetts Institute of Technology, Cambridge, MA 02139, USA

**Keywords:** Posttranscriptional regulation, miRNA dynamics, miRNA stability, metabolic labeling, 5-ethynyluridine

## Abstract

Rates of production and degradation together specify microRNA (miRNA) abundance and dynamics. Here, we used approach-to-equilibrium metabolic labeling to assess these rates for 176 miRNAs in contact-inhibited mouse embryonic fibroblasts (MEFs), 182 miRNAs in dividing MEFs, and 136 miRNAs in mouse embryonic stem cells (mESCs). MicroRNA duplexes, each comprising a mature miRNA and its passenger strand, are produced at rates as fast as 114 ± 49 copies/cell per min, which exceeds rates reported for any mRNAs. These duplexes are rapidly loaded into Argonaute, with <10 min typically required for duplex loading and silencing-complex maturation. Within Argonaute, guide strands have stabilities that vary by 100-fold. Half-lives also vary globally between cell lines, with median values ranging from 6.3 to 34 h in mESCs and contact-inhibited MEFs, respectively. Moreover, relative half-lives for individual miRNAs vary between cell types, implying the influence of cell-specific factors in dictating turnover rate. The apparent influence of miRNA regions most important for targeting, together with the effect of one target on miR-7a-5p accumulation, suggest that targets fulfill this role. Analysis of the tailing and trimming of miRNA 3′ termini showed that the flux was typically greatest through the isoform tailed with a single uridine, although changes in this flux did not correspond to changes in stability, which suggested that the processes of tailing and trimming might be independent from that of decay. Together these results establish a framework for describing the dynamics and regulation of miRNAs throughout their lifecycle.

## Introduction

MicroRNAs (miRNAs) are small RNAs that mediate post-transcriptional gene regulation by targeting mRNAs for degradation and translational repression. In animals, miRNAs typically recognize their mRNA targets through base pairing of the miRNA seed region (miRNA nucleotides 2–8) and sites in mRNA 3′ untranslated regions (UTRs) (Bartel 2018). Each miRNA has many sites throughout the transcriptome, and more than 60% of human protein-coding genes have evolutionarily conserved sites to at least one of the 90 miRNA families that are conserved to fish (Friedman et al. 2009). When knocked out in mice, most miRNA families conserved to fish exhibit abnormal phenotypes, including embryonic or postnatal lethality, infertility, and developmental defects (Bartel 2018). Many miRNAs are regulated both spatially and temporally; the prevalence and conservation of such regulation implies the importance of precise and specific control of miRNA accumulation in each cell type (Houbaviy et al. 2003; Wienholds et al. 2005; Rybak et al. 2008; Thornton and Gregory 2012; Chen and Qin 2015).

The intracellular accumulation of an RNA species is a function of both its production and its destruction, but how the balance between these two processes impinges on miRNA levels is poorly understood. MicroRNA production is a multi-step pathway with opportunities for regulation at each step. Canonical miRNAs are initially transcribed as long, hairpin-containing transcripts known as ‘pri-miRNAs’ (Lee et al. 2002). In animal cells, each miRNA hairpin within the pri-miRNA is cropped in the nucleus by the Microprocessor complex, producing a ‘pre-miRNA’ that is then exported to the cytoplasm and cleaved by Dicer to yield the mature miRNA duplex (Ha and Kim 2014). One strand of this duplex is loaded into an Argonaute (AGO) protein to generate a mature complex capable of targeting genes for repression, whereas the other strand is rapidly degraded (Kawamata and Tomari 2010). Strand selection is typically biased, dictated both by the sequence and the thermodynamic stability of the 5′ ends of each duplex strand (Suzuki et al. 2015; Khvorova et al. 2003; Schwarz et al. 2003), and thus for most miRNA duplexes the strand that is preferentially loaded as the guide strand is readily distinguished from the one that usually acts as the passenger strand. Production of the mature complex can be modulated at each of the steps of biogenesis in a miRNA-specific manner (Lee et al. 2007; Krol et al. 2010; Viswanathan et al. 2008; Heo et al. 2008; Trabucchi et al. 2009; Ma et al. 2010; Han et al. 2007).

In contrast to our understanding of miRNA biogenesis, less is known about miRNA degradation. For example, the extent to which miRNAs degrade through a common pathway characterized by stereotypic changes is unknown. Some mature miRNAs within AGO can have nucleotides added to or removed from their 3′ ends, processes referred to as tailing and trimming, respectively (Ameres and Zamore 2013). Although such modifications can alter stability of some miRNAs, levels of many miRNAs are insensitive to depletion of individual tailing factors (Jones et al. 2009; Katoh et al. 2009; Mansur et al. 2016). Tailing and trimming is also associated with target RNA-directed miRNA degradation (TDMD), a phenomenon in which a highly complementary target site that contains extensive pairing to the 3′ end of a miRNA can trigger degradation of that miRNA (Ameres et al. 2010; Cazalla et al. 2010; Marcinowski et al. 2012; de la Mata et al. 2015; Bitetti et al. 2018; Kleaveland et al. 2018). However, whether trimming and tailing plays a general role in the miRNA decay pathway remains to be determined. Indeed, elimination of tailing by knockout of the Gld2 cytoplasmic poly(A) polymerase has no effect on TDMD-mediated destabilization of miR-7 by the Cyrano long noncoding RNA (lncRNA) in mammalian tissues (Kleaveland et al. 2018).

Questions also remain regarding the rates of miRNA turnover, and the extent to which degradation is regulated in a miRNA-specific manner. Most miRNAs that have been examined are highly stable. In rat hearts miR-208 persists for weeks after its production has ceased, and in several human and mouse cell lines the levels of most miRNAs do not detectably change after up to 24 h of transcriptional inhibition (van Rooij et al. 2007; Bail et al. 2010; Guo et al. 2015). Furthermore, miRNAs persist long after inducible deletion of *Dicer1* in mouse embryonic fibroblasts (MEFs), implying half-lives of over 100 h (Gantier et al. 2011). In contrast, some neuronal-specific miRNAs exhibit more rapid turnover (Krol et al. 2010), and miR-503, a member of the miR-16 extended family proposed to reinforce cell cycle arrest in G0, turns over with a half-life of ~4 h (Rissland et al. 2011). The correlation between short half-life and functions in regulating rapid cell transitions (i.e., response to neuronal stimulation and exit from cell cycle arrest) implies that miRNA half-lives might be selectively regulated to enable dynamic behavior.

Until recently, a challenge facing the field of miRNA degradation has been the lack of a high-throughput but non-disruptive method for assessing turnover rates. Inhibition of transcription dramatically disrupts cellular homeostasis, limiting the duration over which decay can be assessed and calling into question the relevance of the half-lives observed (Bensaude 2011; Sun et al. 2012; Lugowski et al. 2018). Metabolic labeling is less invasive and has long been applied to study mRNA half-lives (Ross 1995; Dölken et al. 2008; Rabani et al. 2011; Schwanhäusser et al. 2011; Duffy et al. 2015) yet has lagged in its application to miRNAs because their short lengths require very efficient and selective pulldown to observe signal above background. Two recently published studies use metabolic labeling with 4- thiouridine to assess rates of miRNA turnover in HEK293T cells and 3T9 mouse fibroblasts, respectively (Duffy et al. 2015; Marzi et al. 2016). However, the former study does not provide half-life values but instead classifies miRNAs as having either fast or slow turnover relative to the bulk population, and the latter study reports that only ~30% of miRNAs have half-lives of greater than 24 h, which seems at odds with results from previous approaches.

Metabolic incorporation of 5-ethynyluridine (5EU) into nascent RNA, followed by attachment of a biotin or fluorescent label through click chemistry acting at the ethynyl group of 5EU, provides a powerful method for isolating or detecting newly synthesized RNA (Jao and Salic 2008; Chan et al. 2018; Kwasnieski et al. 2019). The adaptation of this approach to efficiently isolate ~30-nt fragments of newly synthesized mRNAs (S Eichhorn, T Eisen, D Bartel, unpublished results) suggested that when examining the production and decay dynamics of RNAs as short as miRNAs, 5EU metabolic labeling might offer a favorable alternative to 4-thiouridine labeling. Here, we applied 5EU-labeling to assess the dynamics of miRNA production and decay in contact-inhibited MEFs, dividing MEFs, and mouse embryonic stem cells (mESCs). The results elucidate many previously inaccessible aspects of the miRNA lifecycle, such as the dynamics of duplex loading and the role of trimming and tailing of the miRNA, and lay the foundation for understanding regulation of miRNA degradation and its role in miRNA biology.

## Results

### Reproducible and verifiable high-throughput measurement of miRNA half-lives

Approach-to-equilibrium labeling experiments generate information regarding both the production and degradation rates of the species examined (Neymotin et al. 2014). For such experiments, a label is introduced to cells at time zero, and samples are collected over subsequent time intervals (Fig. 1A). We used this strategy to measure miRNA production and decay in contact-inhibited MEFs, collecting cells 1, 2, 4, 8, 24, 72, and 168 h (1 week) after introducing 5EU to the media. From each of these samples, ~5 μg of total RNA was set aside for small-RNA sequencing, which reported the miRNA profile of the unselected input for each time interval. To the remainder of the total RNA, radiolabeled quantification standards were added (a mixture of three 22–25-nt RNAs that each contained a single 5EU and a 30-nt RNA that contained no 5EU), and 5EU-containing small RNAs were enriched and sequenced (Fig. 1A).

**Figure 1.**
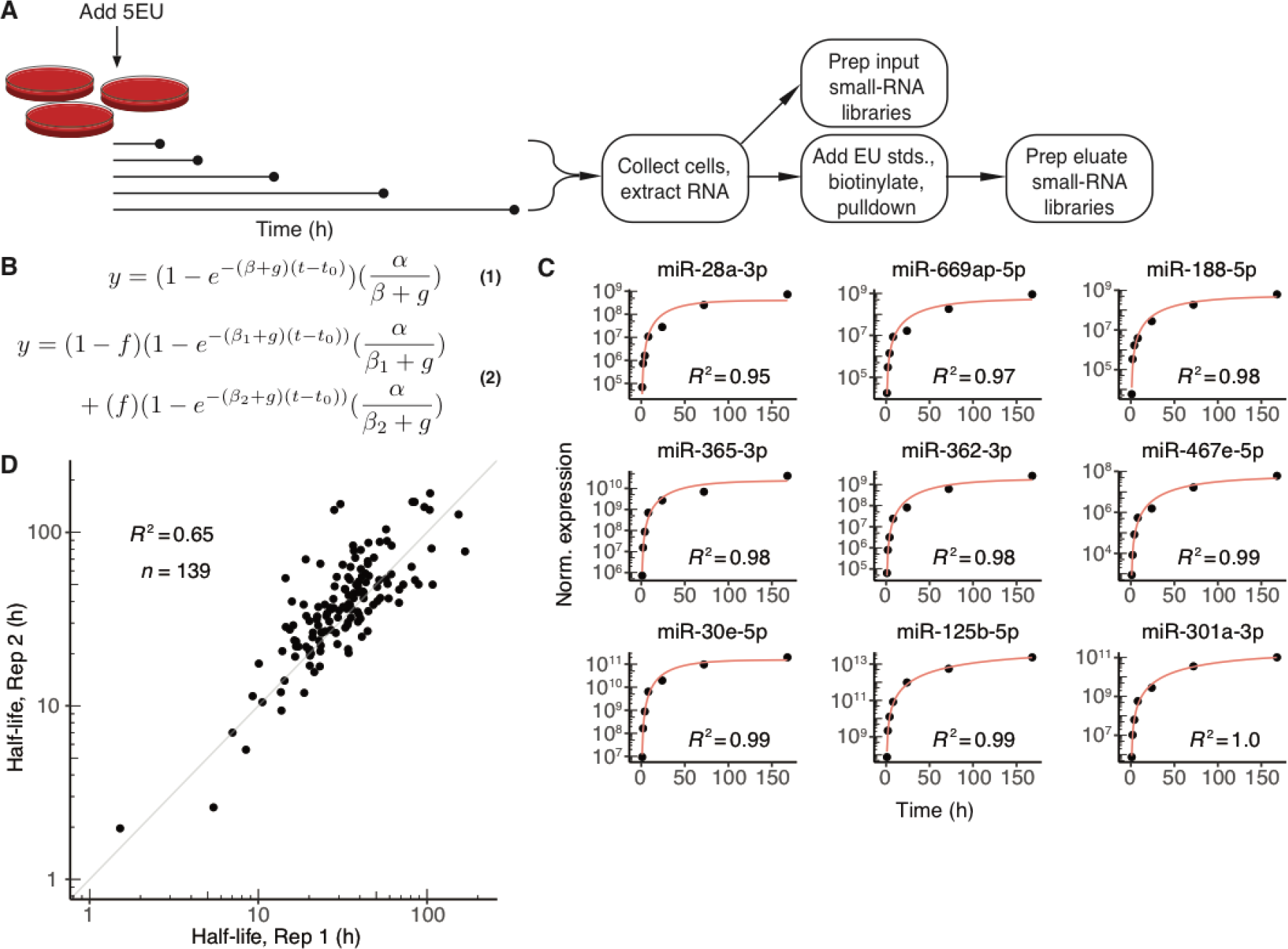
Labeling with 5EU generates reproducible miRNA half-life values. (A) Schematic of the experimental set-up. 5EU is added at time zero, and samples are collected at subsequent time intervals. RNA is extracted from each sample, and a small fraction of this extracted RNA is used to generate small-RNA sequencing libraries. The remainder of the RNA is mixed with quantitative standards (stds.), and molecules with one or more 5EU nucleotides are biotinylated and captured using streptavidin beads. RNA eluted from the beads is prepared for small-RNA sequencing. (B) Equations used for fitting approach-to-equilibrium data. Equation 1 represents the single-exponential model used to fit a single decay rate for guide strands. Equation 2 represents the biexponential model used to fit two decay rates for both passenger and guide strands when assessing passenger-strand half-lives. Fits returned estimates for *α* (production rate), *β*_*n*_ (degradation rate, with *n* distinguishing the two rates of equation 2), *f* (fraction of molecules exhibiting the corresponding degradation rate), and *t*_0_ (time offset). The variable *g* describes the rate of cell doubling and was set manually based on experimental observations. Fits were carried out for each miRNA guide or passenger strand that satisfied an expression cut-off of 60 reads per every million miRNA-mapping reads (i.e., 60 RPM) at each timepoint. (C) Representative fits to data from contact-inhibited MEFs. Data are plotted with black points, and single-exponential fits with a red line. Time-courses representing the 10^th^ through 90^th^ deciles in goodness-of-fit are shown in order (from top left to bottom right) of increasing goodness-of-fit. (D) Reproducibility of half-life measurements for guide strands. Shown is the relationship between values determined independently from each of the two biological replicates from contact-inhibited MEFs.

To assess the efficacy of 5EU labeling and the applicability of approach-to-equilibrium labeling for determining rates of miRNA production and degradation, we examined enrichment of both the 5EU-containing standards and cellular miRNAs over non-labeled species. Monitoring the fate of the internal quantification standards after affinity purification of 5EU-containing species showed that standards with a single 5EU were enriched >150-fold relative to the standard that lacked a 5EU (Supplemental Fig. S1A,B). Levels of cellular miRNAs at each timepoint were also compared to background levels, which were determined by subjecting samples from unlabeled cells to biotinylation and affinity enrichment (Supplemental Fig. S1C). As the duration of 5EU treatment lengthened, yields of pulled-down miRNAs progressively increased over background levels, presumably due to higher proportions of 5EU-containing miRNAs following longer labeling intervals (Supplemental Fig. S1C). As expected, miRNAs with fewer uridines tended to be recovered somewhat less efficiently, but only one uridine-depleted miRNA was poorly recovered in the pull down (Supplemental Fig. S1D). Importantly, treatment for up to one week with 5EU had little effect on cellular miRNA levels (Supplemental Fig. S1E). Together, these observations indicated that metabolic labeling with 5EU could provide an effective method to measure miRNA production and decay, with minimal disruption to miRNA homeostasis.

We fit the results of this approach-to-equilibrium experiment to an exponential function, which generated values for the production and degradation rates (*α* and *β* respectively) of each miRNA guide strand (including passenger strands as well as guide strands) that exceeded our expression threshold (Fig. 1B, equation 1; Supplemental Table S1). In addition to the sequencing data, the exponential model took as input the rate of cell division (*g*), which was experimentally determined for each cell line studied. Data for all miRNAs were fit simultaneously to yield, in addition to miRNA-specific production and degradation rates, a single time offset parameter (*t*_0_) for the experiment that accounted for the lag in the signal due to the time required for 5EU incorporation and miRNA processing. The single-exponential model fit the data well (Fig. 1C), and improvement of fits obtained when using a bi-phasic, biexponential model was insufficient to justify use of this more complex model.

MicroRNA guide strand half-lives and production rates determined from two biological replicates correlated well (Fig. 1D and Supplemental Fig. S1F, Pearson *R*^2^ = 0.65 and 0.92, respectively). This reproducibility allowed data from replicates to be pooled and fit simultaneously (Supplemental Table S1), which generated higher-confidence values for production and degradation rates. The half-life measured for miR-503 in contact-inhibited MEFs using these pooled data, at 6.9 h, agreed well with its previously published half-life in 3T3 cells (Rissland et al. 2011). Together, the analysis of biological replicates and the validation of miR-503 turnover rate confirmed that rates of production and degradation determined from the single-exponential fits to the metabolic-labeling data were both reproducible and accurate.

### Rates of miRNA guide-strand production and degradation

For the guide strands of miRNA duplexes, rates of both production and degradation varied broadly, spanning three and two orders of magnitude, respectively (Fig. 2A,B; Supplemental Table S1). Knowledge of the absolute number of miR-7 molecules in contact-inhibited MEF cells enabled production rates to be converted to absolute values of copies/h per cell (Supplemental Fig. S2A). These analyses revealed that miR-26a-5p, the third most abundant miRNA in these cells, was produced at the fastest rate, with 20 ± 7 (± s.d.) molecules being produced each minute in the average cell. Four additional miRNAs were produced at rates exceeding 10 molecules/min per cell. Production rates of species deriving from genes separated by <10,000 bp were significantly more similar than those of genes separated by >100,000 bp (*p =* 0.00016, Mann-Whitney test). This clustering of similar production rates agreed well with previous estimates of pri-miRNA transcript length (Baskerville and Bartel 2005; Chang et al. 2015), which further supported the accuracy of our production-rate measurements. Rates of production and degradation were not correlated, reflecting independence of the miRNA biogenesis and decay pathways (Supplemental Fig. S2B). Although most guide strands (127/176) had half-lives exceeding 24 h (median half-life, 34 h), nine guide strands had half-lives < 10 h (Fig. 2B). These relatively short-lived species included miR-503, as previously mentioned, as well as miR-7, the miRNA with the shortest measured half-life (Fig. 2B, 1.7 h) and a known substrate of TDMD (Kleaveland et al. 2018).

**Figure 2.**
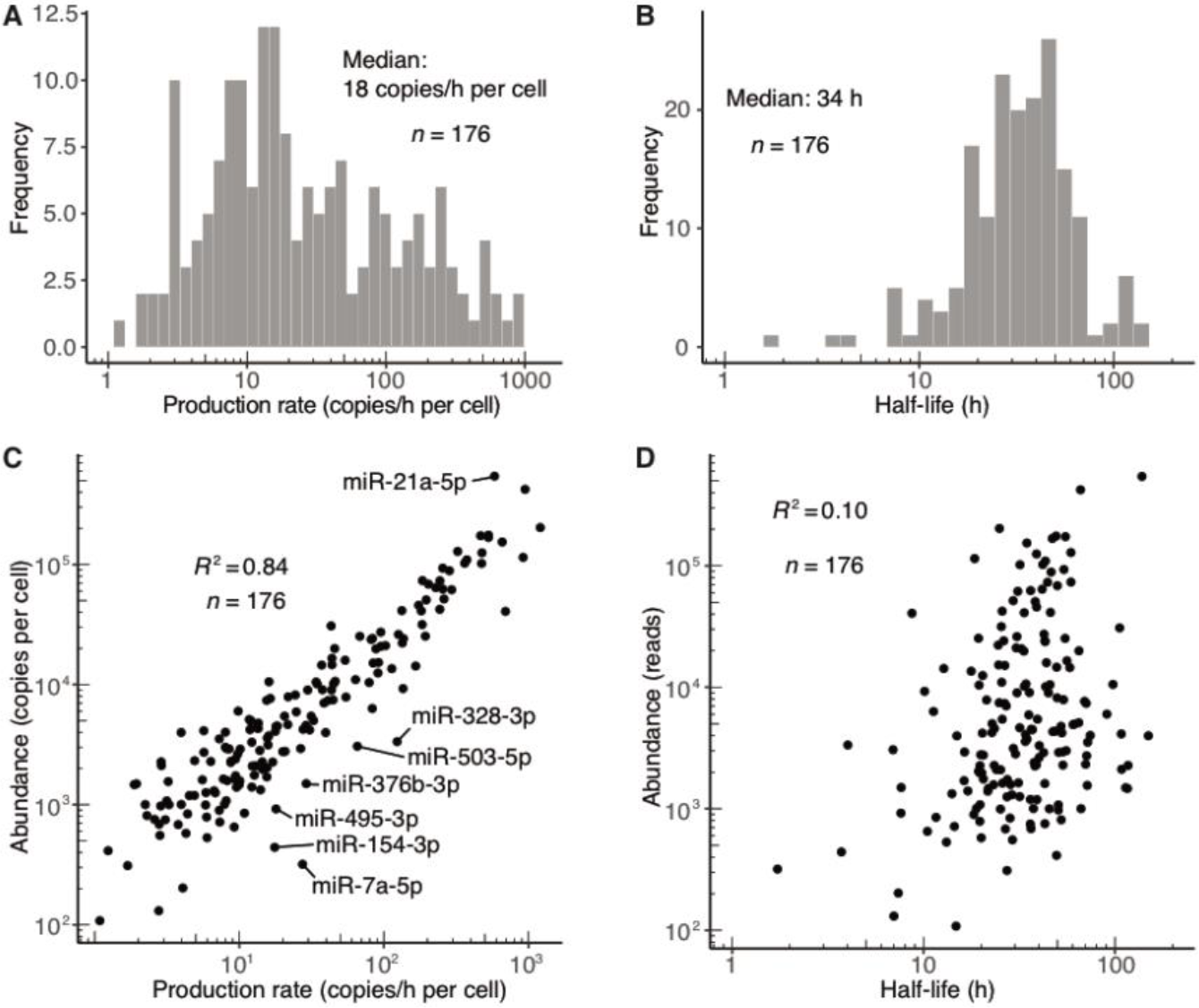
Dynamics of miRNA guide strands in contact-inhibited MEFs. (A) Distribution of production rates for guide strands in contact-inhibited MEFs. (B) Distribution of half-life values for guide strands in contact-inhibited MEFs. (C) Relationship between steady-state abundance (reads in the 1-week input sample) and production rate of guide strands in contact-inhibited MEFs. Outliers that passed the expression threshold (production rate >5 copies/h per cell) and differed significantly from the best linear fit (|z-score| > 2) are labeled. (D) Relationship between steady-state abundance (reads in the 1-week input sample) and half-life of guide strands in contact-inhibited MEFs.

Production rate correlated well with miRNA abundance (Fig. 2C, Pearson *R*^2^ = 0.84), suggesting that regulated production is the primary cause of differing miRNA levels. Nevertheless, as exemplified by a handful of miRNAs with unusually short half-lives that included both miR-7a-5p and miR-503-5p, some miRNAs were produced at rates substantially higher than would have been expected for their overall abundances (Fig. 2C). Moreover, although the correlation between half-life and abundance was low (Pearson *R*^2^ = 0.10), it was nonetheless significant (P < 10^−4^), and the eight miRNAs with the fastest turnover each had below-median abundance (Fig. 2D). These results indicated that differential stability can substantially impact miRNA accumulation.

### Rates of duplex loading and passenger-strand degradation

Compared to miRNA guide strands, passenger strands are thought to have a more ephemeral existence, starting the moment of Dicer-catalyzed duplex production and lasting through the process of duplex loading and silencing complex maturation, which culminates with either slicing of the passenger strand or its release as a single-stranded species vulnerable to rapid degradation. As such, monitoring the half-life of the passenger strand would be expected to provide a window into the rate of duplex loading and silencing-complex maturation. Complicating this picture, however, strands that most often acted as passenger strands, much more so than guide strands, exhibited bi-phasic approach to equilibrium, shooting up rapidly at early time intervals and then increasing at a slower rate (Fig. 3A). This behavior indicated a mixture of two populations, a major one that was rapidly degraded and a minor one that was more stable. Such behavior would be expected if loading into AGO was not perfectly consistent, and strands that were normally passenger strands were occasionally loaded into AGO as guide strands and thereby stabilized. Of course, this explanation implied that the strand designated as the guide strand of these duplexes occasionally failed to load into AGO, and raised the question of why these guide strands also did not exhibit a bi-phasic approach to equilibrium. The answer to this question lies with the fact that for strands that most often acted as guide strands the minor population was the one that was rapidly degraded, and rapidly degraded minor populations have imperceptible influence on curves describing approach to equilibrium as well as the half-life values inferred from these curves, as demonstrated from simulations (Supplemental Fig. S3A). In contrast, for strands that most often acted as passenger strands the minor population was much more stable than the major one and thus could represent a sizeable fraction of the accumulating RNA. The minor population could thereby influence the shape of the curve to inflate the apparent half-life of the passenger strand such that its value did not accurately reflect the rapid rate of turnover of the majority of the passenger species (represented by the initial burst in accumulation) (Supplemental Fig. S3A).

**Figure 3.**
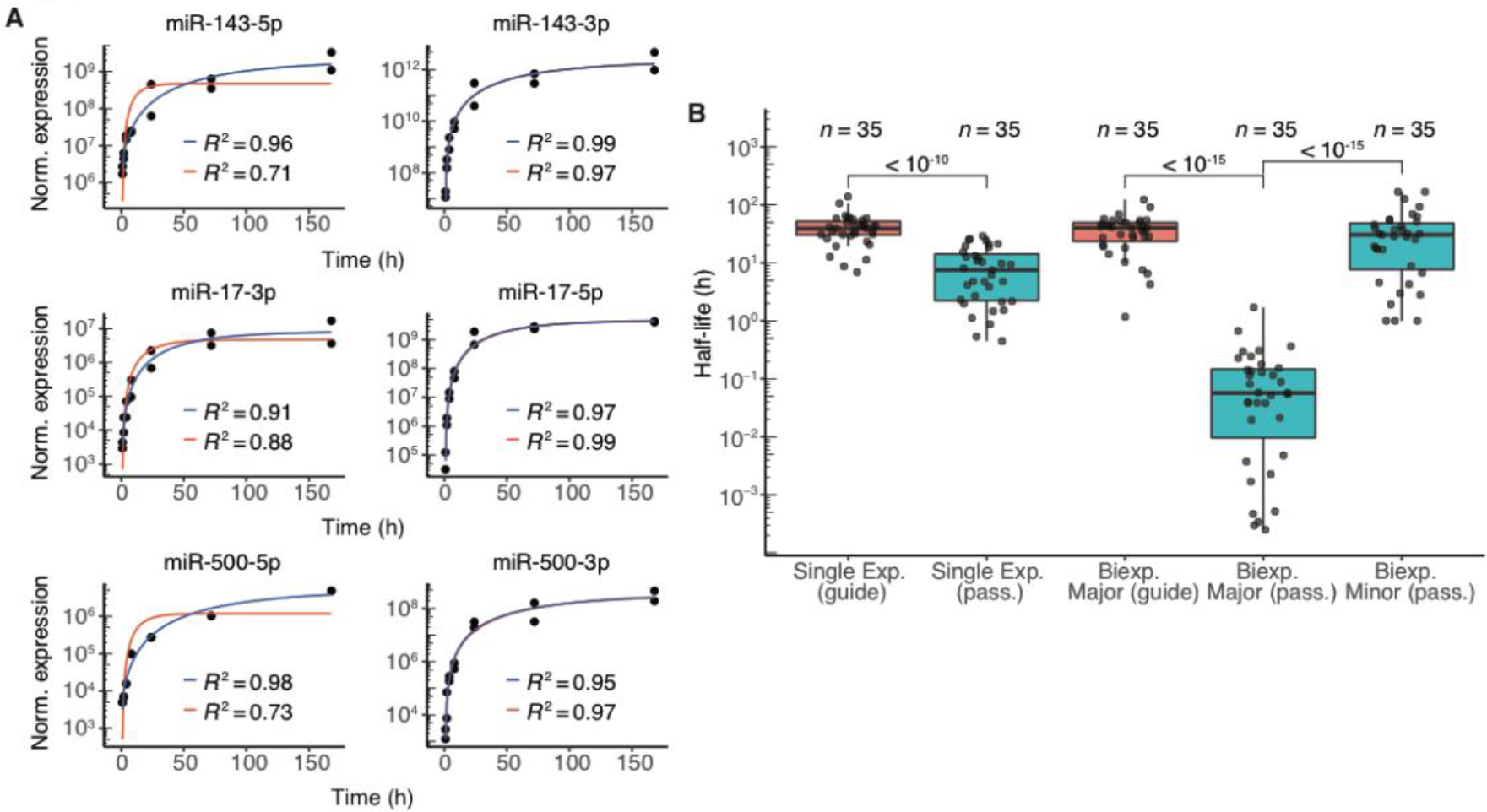
Dynamics of passenger strands in contact-inhibited MEFs. (A) Examples of fits for passenger strands and corresponding guide strands (left and right, respectively). Single- and biexponential fits are indicated (red and blue lines, respectively) with corresponding *R*^2^ values. (B) Half-life distributions for guide and passenger strands obtained using either the single-exponential (single exp.) or biexponential (biexp.) fitting methods (box, quartiles; whiskers, at most 1.5 times the inter-quartile range). The major half-life was the one that corresponded to the majority of the specified strand (i.e., for the passenger strand, the molecules not incorporated into the mature silencing complex, and for the guide strand, the molecules incorporated into the mature complex), whereas the minor half-life was the one that corresponded to the minority of the specified strand. P values (two-sample *t* test) are shown for the indicated comparisons.

These insights indicated that a biexponential model for the approach to equilibrium, which included rates for both the minor and major populations (Fig. 1B, equation 2), was more appropriate for fitting the data of strands that usually acted as passenger strands. However, simulations revealed that fits to the biexponential model were insufficiently constrained when applied to each passenger strand independently; accurate determination of parameters required passenger-and-guide pairs instead to be fit simultaneously. This simultaneous fitting of passenger–guide pairs reduced the number of fitted variables by specifying a single production rate for both strands, which was a reasonable assumption when considering that both strands were produced simultaneously from the same molecule. The simultaneous fitting also constrained the fractions of passenger and guide species loaded into AGO to add up to 1.0, a reasonable assumption from the perspective that only one strand of a duplex can be loaded as the guide (although this constraint did not allow for the possibility that duplex molecules might be degraded before either strand is loaded, see discussion). Simultaneous fitting of parameters for passenger–guide pairs enabled accurate determination of major and minor half-lives of simulated passenger species as well as major half-lives of simulated guide species (Supplemental Fig. S3B,C).

This model was then applied to 35 passenger–guide pairs in the contact-inhibited–MEF datasets for which both strands surpassed our expression threshold but exhibited greater than 5-fold differences in steady-state accumulation (Supplemental Table S2). For guide species deriving from each of these 34 pairs, the major half-lives resembled half-lives obtained from the single-exponential fits (Supplemental Fig. S3D). (For the exceptions in which half-lives fit by the two methods deviated, the biexponential model typically predicted shorter half-lives, which was attributable to the difficulty of accurately fitting the minor half-life of these guide strands, which in turn impacted the guide strand major half-life. Thus, for guide strands, half-lives from the single-exponential fits were considered more accurate and used for subsequent analyses.) The passenger-strand major half-lives, however, were significantly lower than those obtained from the single-exponential fits (*p* < 10^−15^, t-test) and almost three orders of magnitude below guide-strand major half-lives (Fig. 3B, Supplemental Table S2). Moreover, the distribution of passenger-strand minor half-lives did not significantly differ from that of the guide-strand major half-lives (Fig. 3B, *p* = 0.16, t-test), which implied that strands that normally act as passengers, when occasionally loaded into AGO, generally behave no differently than do strands that normally act as guides.

The major half-lives of the passenger strands, interpreted as the half-lives of these strands when they failed to load into AGO, provided insight into the rates of duplex loading and silencing-complex maturation in contact-inhibited MEF cells. The median major half-life of passenger strands was 0.057 h (Fig. 3B), which suggested that once produced, about 5 min is typically required for the duplex to be loaded into AGO and the passenger strand to be either expelled (and subsequently degraded) or sliced.

Moreover, the broad distribution of passenger-strand major half-lives, which ranged from < 0.01 to 1 h, implied that the rates of duplex loading and complex maturation, varies substantially for different miRNA duplexes.

### Sequence differences and other features that mediate differences in guide half-lives

To address what might specify differences in guide half-lives, we assessed whether species with similar sequences had similar half-lives. MicroRNAs with the same seed region target largely overlapping sets of mRNAs and are classified as members of the same family (Bartel 2009). To assess whether these seed nucleotides also help to specify half-life, we compared differences in half-life for pairs of family members to those for random, non-family miRNAs pairs. Half-lives were not significantly more similar for family members (Fig. 4A), which indicated that the seed sequence alone does not specify half-life. Because family members target essentially the same mRNAs, these results do not support a proposal that the sheer abundance of target mRNAs can influence miRNA stability (Chatterjee et al. 2011).

**Figure 4.**
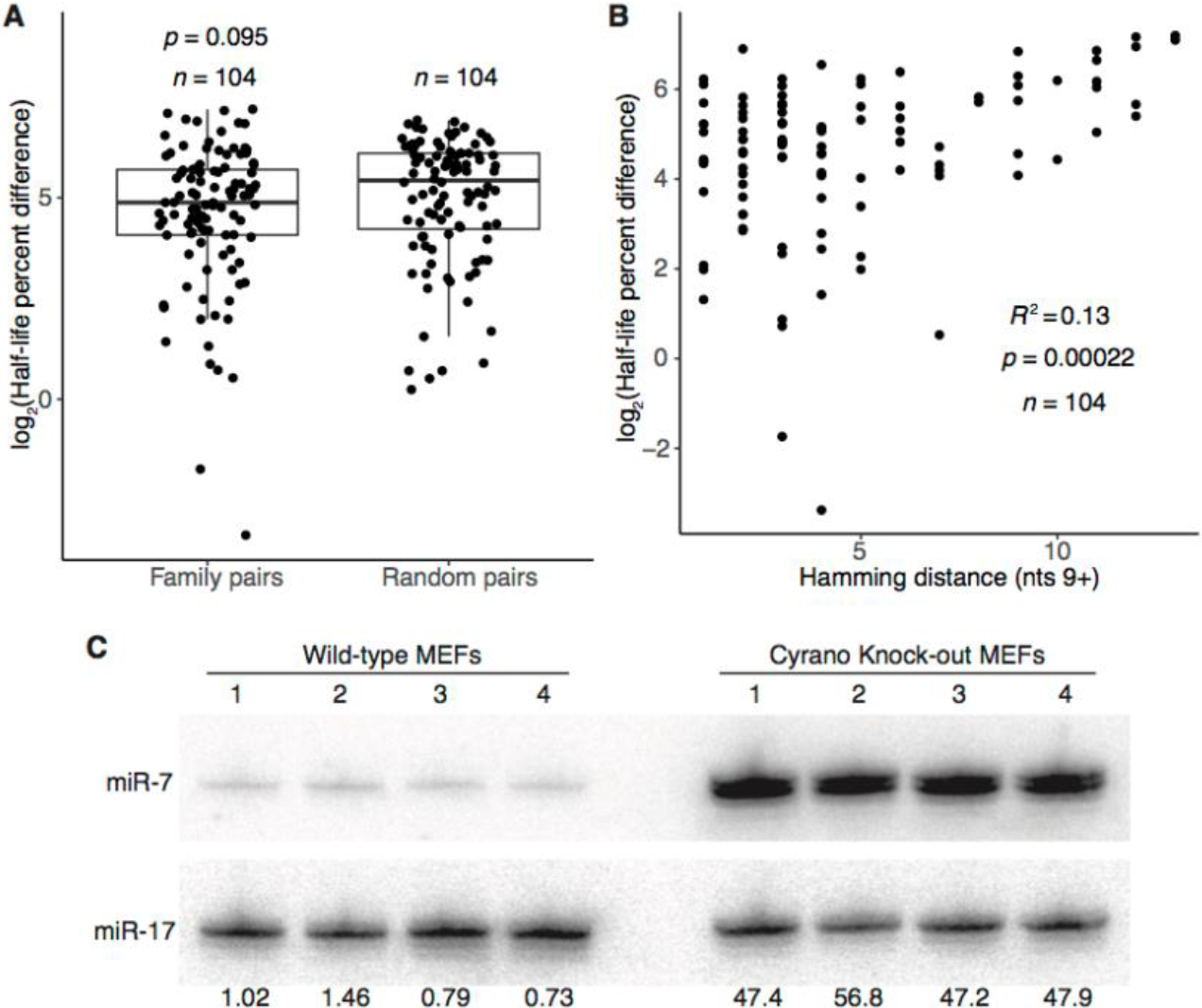
Features associated with rapid miRNA turnover. (A) Differences in half-life for pairs of family members or randomly paired guides. For each pair-wise comparison of all members within each miRNA family, half-life percent differences were computed as the absolute value of the difference between half-lives divided by the mean of the two half-lives, and the log_2_-transformed percent-difference values of all such comparisons in contact-inhibited MEFs are plotted (box, quartiles; whiskers, at most 1.5 times the inter-quartile range). Twenty-five equally sized cohorts of randomly paired guides were analyzed in the same way, with the cohort shown for comparison being the one with the median value of significance (*p* value, two-sample *t* test). (B) Relationship between half-life difference and the number of non-identical nucleotides (Hamming distance) when comparing pairs of family members. Only nucleotides 3′ of the seed region were considered for the Hamming-distance measurement. Significance of the correlation was determined by a t-test. (C) The effect of Cyrano on steady-state miR-7 levels in contact-inhibited MEFs. Shown is a scan of a northern blot analyzing RNA from wild-type and Cyrano-knockout contact-inhibited MEFs. For each cell line, four replicates were analyzed, sequentially probing for miR-17, which served as a loading control, and then for miR-7. The fold-difference in miR-7 relative to the average miR-7 level in wild-type MEFs is shown below each lane.

The failure of the seed region alone to specify half-life indicated that the remainder of the miRNA plays a role in dictating turnover. Indeed, when comparing pairs of family members, sequence similarity outside of the seed region, as measured by Hamming distance, significantly correlated with more similar rates of turnover (Fig. 4B, *R*^2^ = 0.13). This observation did not extend to pairs of non-family-member miRNAs (Supplemental Fig. S4A, *R*^2^ = 0.0033), which indicated that although the seed region alone is not sufficient to specify half-life, it does play a role in half-life specification. To identify portions of the non-seed region that might be especially important in driving half-life similarities, we repeated the correlation analysis for all 4-nt non-seed segments of family-member pairs. Although hamming distance and half-life similarity correlated for most 4-nt segments, the correlation peaked for the segment spanning miRNA nucleotides 13–16 (Supplemental Fig. S4B). This region of the miRNA has the greatest propensity to engage in supplemental pairing to targets, and approximately 5% of conserved seed-matched sites exhibit conserved pairing to this segment (Grimson et al. 2007; Friedman et al. 2009). Together, these observations indicate that regions that preferentially engage in target pairing also help to specify half-life.

In an attempt to identify the features that specify half-life, we searched for common motifs or traits among guide strands that rapidly turned over, considering overall nucleotide content, position-specific nucleotide content, overall dinucleotide or trinucleotide motifs, position-specific *k*-mers 1–4 nt in length, and predicted pairing stabilities of the seed and supplemental regions. However, no consistent or striking enrichments or relationships were observed (Supplemental Fig. S4C,D, data not shown). Moreover, no correlation was observed between half-life and either the abundance of all seed-matched, high-confidence predicted targets or the abundance of all such targets with extended supplemental pairing (Supplemental Fig. S4E). Our inability to identify a general feature specifying rapid miRNA turnover implied either that the causative factor had not been properly probed or that unique factors specified turnover of the different rapidly degraded species.

Of the guide strands expressed in contact-inhibited MEF cells, the least stable was miR-7, with a half-life of only 1.7 h measured in our high-throughput analysis. This miRNA pairs extensively to a site within the Cyrano lncRNA, which can promote its efficient destruction through TDMD (Kleaveland et al. 2018). To assess whether Cyrano might be influencing miR-7 stability in contact-inhibited MEFs, we measured miR-7 steady-state levels in both wild-type and Cyrano-knockout cells. In the absence of Cyrano, the miR-7 level increased ~50-fold (Fig. 4C), suggesting that the presence of one highly complementary target can dramatically alter the stability of a miRNA. This strong influence of a single target might explain why no correlation was observed between half-life and the overall abundance of predicted targets with supplemental pairing. When instead target abundance of the top *n* species with the most supplemental pairing to each miRNA was assessed for *n* from 1 to 50, however, no comparison had a Pearson *R*^2^ > 0.03. This result suggested either that we have not yet identified the Cyrano-like targets for the other rapidly turned-over guide strands, or that miR-7 is an outlier in having its half-life largely shaped by its interaction with a single highly complementary target.

### Different miRNA dynamics in dividing MEFs and mECSs

To get a sense of the cell-specific nature of miRNA dynamics, we extended our high-throughput, metabolic-labeling approach to both dividing MEFs and mESCs. As observed for contact-inhibited MEFs, production rates varied broadly for individual miRNAs within each cell line—spanning three or more orders of magnitude (Supplemental Fig. S5A,B). Comparing rates of production across the different experiments reflected the degree of similarity between the cell lines compared, with contact-inhibited MEFs correlating well with dividing MEFs but hardly at all with mESCs (Supplemental Fig. S5C,D, *R*^2^ = 0.78 and 0.040, respectively). To compare absolute rates of production in the two MEF lines, the number of miR-7 molecules in each dividing MEF cell was determined and used to calculate absolute production rates in dividing MEFs (Supplemental Fig. S2C). These analyses revealed that miR-21a-5p, the most abundant miRNA in these cells, was produced at a rate of 114 ± 49 molecules/min per cell, and that seven other miRNAs were produced at rates > 10 molecules/min per cell. Although this maximum production rate was ~6-fold greater than what was observed in contact-inhibited MEFs, similar numbers of miRNAs had production rates exceeding 10 molecules/min per cell in the two cell states.

Half-lives in dividing MEFs tended to be slightly shorter than in contact-inhibited MEFs (median half-lives of 25 h and 34 h, respectively; distributions significantly different, *p =* 9 × 10^−5^, Kolmogorov–Smirnov test), and half-lives in mESCs tended to be much shorter (median, 6.0 h; *p <* 10^−15^) (Fig. 5A,B). In addition to an overall shift in miRNA turnover dynamics, half-lives for individual miRNA guide strands in different cell states or lines exhibited low correlation (Fig. 5C,D). These correlations were noticeably lower than those between replicates within a given cell line (Fig. 1D, Supplemental Fig. S5E), which indicated that different cell lines, and to some extent, different cell states, can have significant differences in individual miRNA turnover over and above differences observed in bulk turnover. A notable exception to this cell-type-specific behavior was miR-7, which was among the least stable guide RNAs in all three datasets. Nonetheless, the observation that individual miRNAs exhibited cell-state– and cell-type–specific posttranscriptional regulation implied differential expression of miRNA-extrinsic factors that influence miRNA degradation.

**Figure 5.**
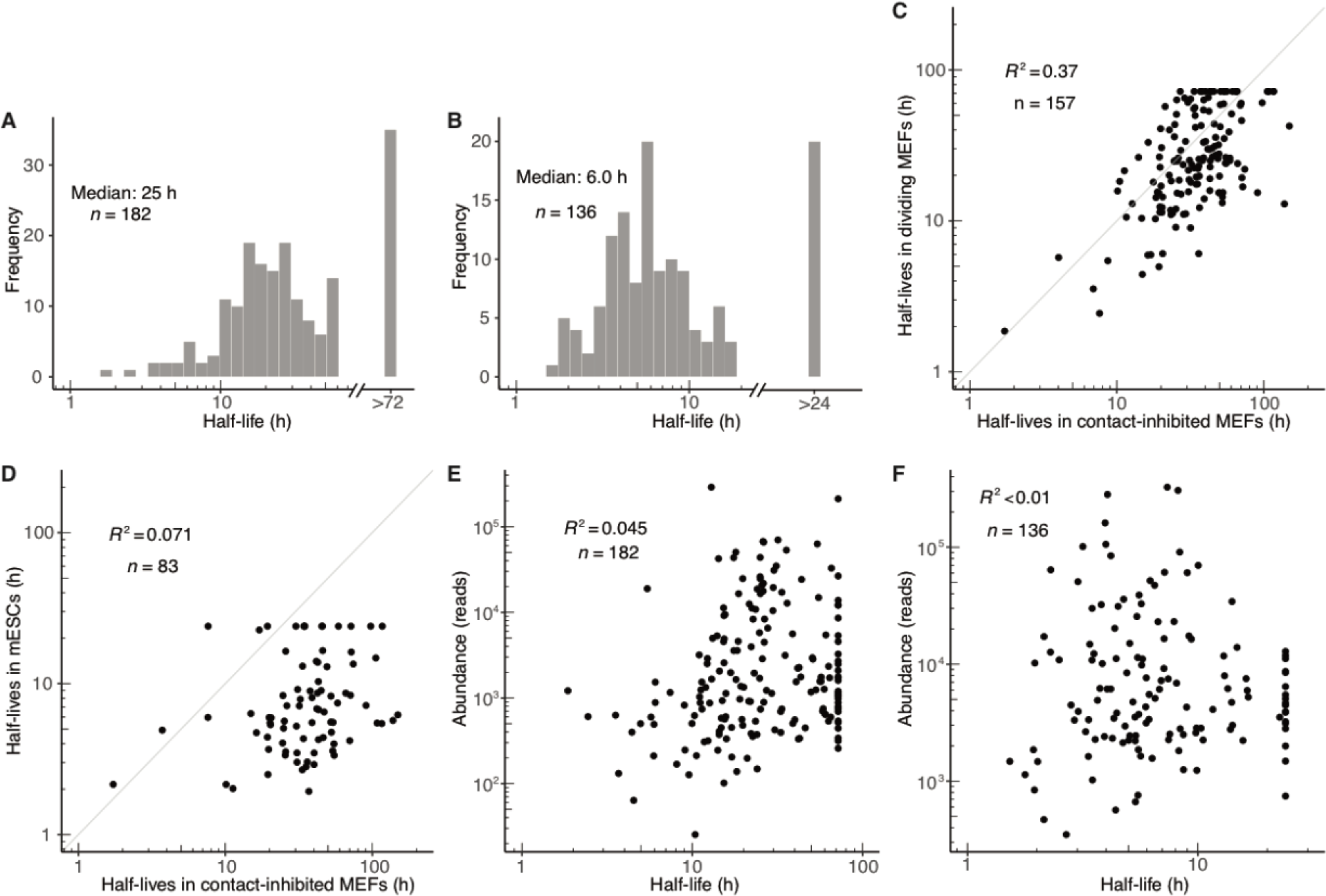
Dynamics of miRNA guide strands in dividing MEFs and mESCs. (A–B) Half-life distributions for guide strands in dividing MEFs (A) and mESCs (B). Half-lives were capped at the duration of the longest time interval (72 h and 24 h in dividing MEFs and mESCs, respectively), as modeling indicated that longer half-lives would be fit less accurately. (C–D) Guide-strand half-lives in dividing MEFs (C) and mESCs (D) as a function of those half-lives in contact-inhibited MEFs. (E–F) Relationship between steady-state abundance and half-life in dividing MEFs (E) and mESCs (F).

Reasoning that global differences in miRNA turnover dynamics might reflect differences in AGO turnover rates in the different cell states and cell lines, we used pulsed SILAC (Schwanhäusser et al. 2011) to measure the half-life of AGO2 in these different cellular contexts. We focused on AGO2 for two reasons: as the most highly expressed AGO family member in each of these contexts, it was presumably the one that was associated with most miRNA molecules of these cells. In addition, AGO2 was the AGO family member that could be reliably queried in all three cellular contexts using our approach. In contact-inhibited MEFs the AGO2 half-life was 57.3 ± 1.6 h (± s.d.), whereas in dividing MEFs the AGO2 half-life decreased to 40.9 ± 2.0 h (Supplemental Fig. S5F). For comparison, we computed a single representative half-life for all miRNA guide strands in these cells. In both contact-inhibited MEFs and dividing MEFs these representative half-lives, of 42.5 h and 19.8 h, respectively, were substantially lower than the measured AGO2 half-lives. In mESCs the AGO2 half-life was difficult to determine because the signal from protein degradation could not be distinguished above the signal due to dilution caused by cell division (Supplemental Fig. S5F). Nonetheless, a lower limit for AGO2 half-life was 100 h (estimated based on the ability to determine half-lives up to this value with this methodology for other proteins in mESCs), which was much higher than the single representative half-life of 6.0 h calculated for miRNA guide strands in mESCs. Thus, decreased bulk AGO stability failed to explain the general changes in miRNA turnover dynamics observed between cell states and cell types, which indicated that differences in other factors must be contributing to these changes.

To assess the generality of the principles of miRNA dynamics determined from analyses of the contact-inhibited–MEFs dataset, we applied the same sets of analyses to the dividing-MEF and mESC datasets. Abundance of miRNAs was again highly correlated with miRNA production rate and lowly correlated with miRNA degradation rate, with miR-7 in dividing MEFs being the most notable example of a miRNA for which accumulation was substantially affected by degradation rate (Fig. 5E,F, Supplemental Fig. S5G,H). Use of the biexponential model to fit passenger-strand half-lives in dividing MEFs and mESCs corroborated the stability change of over two orders of magnitude attributable to loading into AGO (Supplemental Fig. S6A,B). The median duration of duplex loading followed by passenger-strand decay in dividing MEFs and mESCs was estimated to be 29 and 2.8 min, respectively, which was similar to the 5 min value estimated in contact-inhibited MEFs (Supplemental Fig. S6A,B). Analysis of miRNA family members in dividing MEFs supported the conclusion that regions preferentially involved in target recognition play a greater role in specifying half-life (Supplemental Fig. S6C,D). The analogous analyses for guide RNAs in mESCs was less conclusive, perhaps due to lower numbers of family members and therefore less analytical power (Supplemental Fig. S6E,F). More in-depth analyses of the features that specify miRNA turnover echoed those carried out in contact-inhibited MEFs, with no motifs or commonalities discernible for miRNAs in either dividing MEFs or mESCs.

Acquisition of half-life measurements for miRNAs in dividing MEFs provided the most suitable dataset for comparing our results to those that have been previously reported, which were for miRNAs in dividing mouse 3T9 fibroblasts (Marzi et al. 2016). Half-lives for guide strands determined in both datasets had no correlation (Supplemental Fig. S7A). This result, which was unexpected given the highly correlated mRNA expression observed in the two cell lines (Supplemental Fig. S7B, *R*^2^ = 0.87), could stem from the different experimental protocols used to determine half-lives. Approach-to-equilibrium labeling with 5EU coupled to time-course normalization with quantitative EU-containing standards should in theory improve the sensitivity and accuracy of metabolic labeling as applied to miRNAs, as biotinylation of 4-thiouridine with HPDP-biotin is known to be inefficient (Duffy et al. 2015), and sample normalization with quantitative standards is more robust than normalization against a selected miRNA.

### Rates of trimming and tailing, and their relationship with miRNA stability

After loading into AGO, the 3′ ends of mature miRNAs can change through the action of tailing or trimming. These alternative 3′-end isoforms accumulate over time in a manner that reflects their rates of production, their rates of conversion to other isoforms, and any changes in stability that the modified 3′ end might impart. Our analyses showed that for most miRNAs, the initial mature species remained the most abundant throughout the time course, but for a few, the level of a tailed or trimmed isoform either approached or surpassed that of the initial isoform, as exemplified by miR-23a-3p and miR-674-5p (Fig. 6A). Classes of isoforms exhibited different behaviors for different miRNAs; although all miR-674-5p isoforms accumulated to greater abundance than the mature isoform, only the trimmed isoform did so for miR-23a-3p.

**Figure 6.**
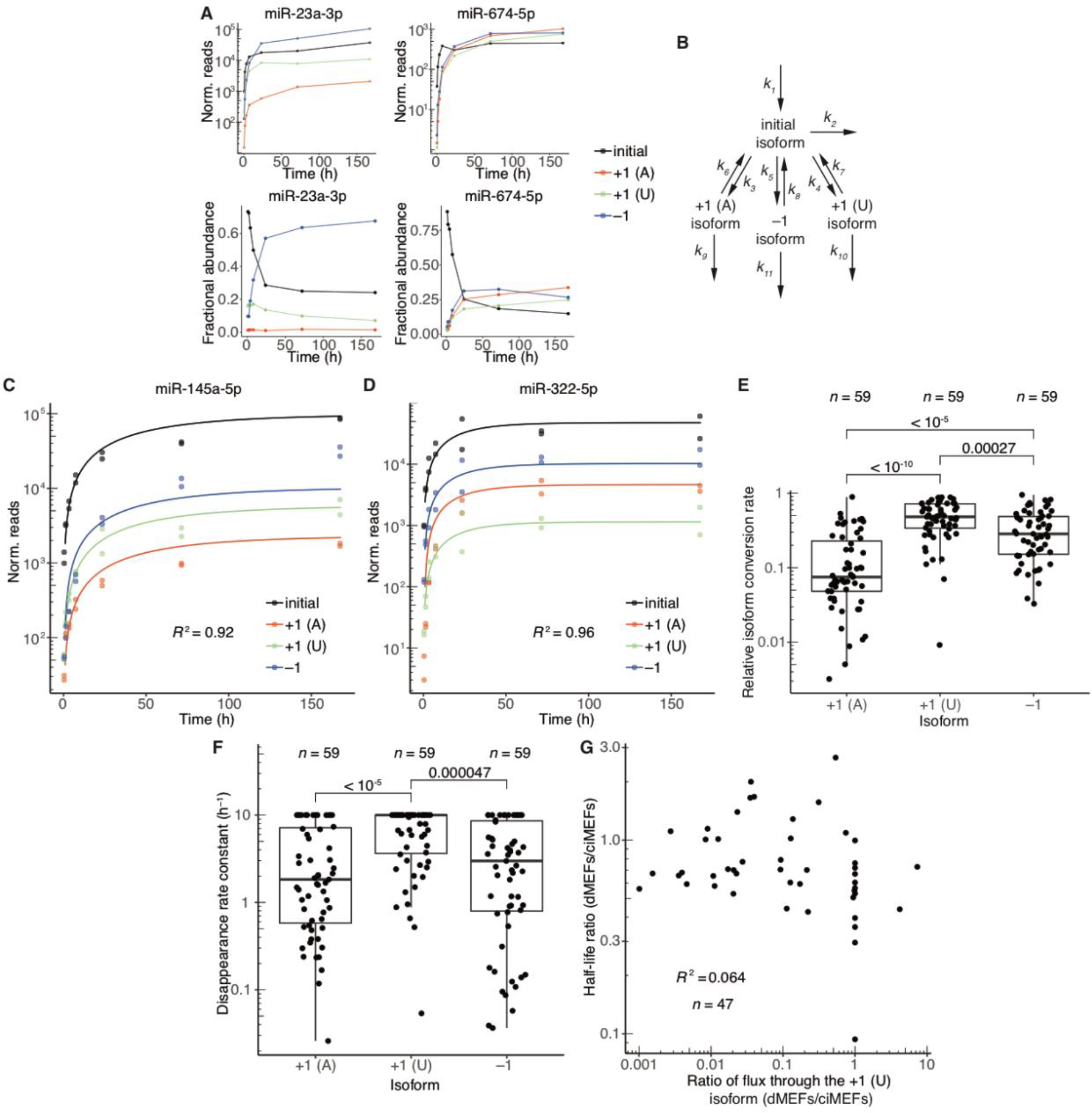
Dynamics of 3′-end isoforms. (A) Isoform accumulation for miR-23a-3p and miR-674-5p, two guide RNAs for which accumulation of derivatives of the initial isoform overtook that of the initial isoform. Top: Absolute accumulation of the initial isoform (black) and its major derivatives (key), in which it is either trimmed by 1 nt (−1, blue) or tailed with either a single A (+1 (A), red) or a single U (+1 (U), green). Lines connect the mean values for each timepoint and do not represent fits to the data. Bottom: Fractional abundance of the initial isoform and its major derivatives. (B) Schematic depicting the isoform-conversion model that was fit to the data and used for simulations. Rate constants extracted from the fit are labeled (*k*_*n*_). Isoform nomenclature is as in (A). (C–D) Representative examples of fits to isoform dynamics for miRNAs in contact-inhibited MEFs. Shown is the plot from the middle of the lowest quartile (miR-145a-5p, *R*^2^ = 0.92) and the plot from the middle of the highest quartile (miR-322-5p, *R*^2^ = 0.96) of goodness-of-fit (*R*^2^), when fitting data from both replicates. (E) Relative rates of conversion to the +1 (A), +1 (U), and −1 isoforms in contact-inhibited MEFs. Rate constants of conversion to each isoform (*k*_3_, *k*_4_, and *k*_5_ for +1 (A), +1 (U), and −1, respectively) were normalized to the summed conversion rate constant (*k*_3_ + *k*_4_ + *k*_5_) to generate the relative rates of conversion. Simulations indicated that conversion rate constants >1 h^−1^ could not be accurately fit, and thus conversion rate constants were capped at this value. Significance was evaluated with a Mann–Whitney test. (F) Rate constants for disappearance (*k*_6_+ *k*_9_, *k*_7_+ *k*_10_, or *k*_8_+ *k*_11_) of the +1 (A), +1 (U), and −1 isoforms, respectively, in contact-inhibited MEFs. Simulations indicated that disappearance rate constants >10 h^−1^ could not be accurately fit, and thus disappearance rate constants were capped at this value. Significance was evaluated with a Mann–Whitney test, and all significant comparisons are indicated. (G) Correlation between half-life ratios (dividing values from dividing MEFs by those from contact-inhibited MEFs) and ratios of the flux through the +1 (U) isoform (again dividing values from dividing MEFs by those from contact-inhibited MEFs). Dividing MEFs denoted as dMEFs, contact-inhibited MEFs denoted as ciMEFs.

To assess the effect of trimming and tailing in a more global manner, and to tease apart the contributions of production and degradation rates to isoform accumulation, we generated a model to describe mature miRNA production and isoform conversion (Fig. 6B). The model fits rate constants of production and disappearance of the mature miRNA (*k*_1_ and *k*_2_, respectively), rate constants of conversion to selected trimmed and tailed species (*k*_3_, *k*_4_, and *k*_5_), as well as rate constants of back-conversion and disappearance for each of the selected species (*k*_6_, *k*_7_, *k*_8_, *k*_9_, *k*_10_, and *k*_11_). This model encompassed all major species that accumulated over the time course but excluded minor species that were detected but never reached a level to be accurately modeled. Thus, each disappearance rate constant accounted for not only decay but also conversion to any isoform not specified by the model. The model was found to accurately fit simulated datasets that spanned a wide array of production, conversion, and degradation rate constants (Supplemental Fig. S8A). These simulations showed that although not all rate constants modeled could be fit accurately with the limited amount of data provided to the model, the model successfully determined four individual rate constants (*k*_1_, *k*_3_, *k*_4_, and *k*_5_) and the sums of three pairs of rate constants (*k*_6_ + *k*_9_, *k*_7_ + *k*_10_, *k*_8_ + *k*_11_).

Application of the model to the contact-inhibited MEF dataset required annotation of the mature species initially produced for each miRNA. These species were annotated using the 1 h timepoint; any species that represented more than 20% of the reads from an arm of a miRNA hairpin were considered Dicer products. This threshold seemed reasonable given the assumption that modified species would not be able to accumulate to such levels after such a short labeling time, especially when accounting for the observed lag time of 36 min (Fig. 1B, *t*_0_ of equation 1). Mature species from 3p arms annotated in this manner were compared with published pre-miRNA sequences of Drosha products in human cells lines (Kim 2017). For 77% of the miRNAs present in both datasets, the 3′ terminus of our inferred Dicer product matched the 3′ terminus of the previously reported Drosha products. The annotations that differed presumably resulted from either modification of the pre-miRNA 3′ terminus prior to Dicer cleavage or species-specific differences in Drosha processing.

We used these annotations and the model to fit rate constants for mature-miRNA production, mature-miRNA conversion to isoforms, and isoform disappearance (which was a combined rate constant describing isoform degradation, isoform conversion to further-modified isoforms, and isoform back-conversion to the mature species) for all miRNAs in the contact-inhibited MEF dataset that passed a threshold requirement of at least 1 read at all timepoints for all isoforms. The model fit the data well, and as expected, it predicted rates of mature-isoform production that were highly correlated with those determined by the single-exponential fits (Fig. 6C,D, Supplemental Fig. S8B,C). Relative rates of conversion to each isoform for individual miRNAs spanned over two orders of magnitude, indicating a high degree of variation in the preferred modification trajectory for different miRNAs (Fig. 6E). As a general trend, however, trimming was faster than A tailing but slower than U tailing (Fig. 6E). Rate constants of disappearance predicted by the model also spanned up to two orders of magnitude for some isoforms, indicating differential responses for individual miRNAs to modifications (Fig. 6F). Disappearance rate constants were largest for U-tailed species, indicating that species containing this modification were destabilized relative to species containing other modifications in contact-inhibited MEFs (Fig. 6F). Rate constants of isoform conversion and isoform disappearance for a given isoform were correlated; miRNAs with relatively high rate constants for conversion to the A-tailed species also had relatively high rate constants of disappearance for this species (Supplemental Fig. S8D). Correlation of these parameters was not observed in fits to simulated data, and thus these correlations did not appear to be an artifact of the model but rather could represent a real biological phenomenon, such as enzyme processivity.

To assess the generalizability of these observations across multiple cell states, we fit the isoform model to data from the dividing MEFs. The parameters fit to these data revealed that global trends in isoform dynamics resembled those observed for contact-inhibited MEFs; in particular, conversion to the U-tailed isoform generally proceeded with the fastest rates, and these U-tailed species were generally less stable than were the other isoforms (Supplemental Fig. S8E,F). At the level of individual miRNAs, rate constants of isoform disappearance were correlated between the two datasets but rate constants of isoform conversion were less so (Supplemental Fig. S8G,H). Both the conversion and disappearance rate constants tended to be somewhat decreased in dividing MEFs as compared to contact-inhibited MEFs (Supplemental Fig. S8G,8H). Correlations between changes in isoform dynamics and changes in half-lives for miRNAs in these two cell states were examined to assess the relationship between flux through isoforms and miRNA turnover. The strongest correlation was observed between the change in flux through the +1 (U) isoform and the change in half-life (*R*^2^ = 0.064, Fig. 6G). Reasoning that a stronger correlation might have been masked by the inability to accurately determine rate constants above certain values (as determined from the isoform dynamics simulations), we repeated these analyses after removing all species with capped rate constants but still observed only very low *R*^2^ values. These low coefficients of determination, in addition to the observations that general flux through isoforms was decreased whereas miRNA turnover rate was increased in dividing MEFs, implied substantial independence between the miRNA modification and degradation pathways.

## Discussion

Our global analyses of miRNA metabolism with 5EU provided half-life measurements for the guide strands of 201 miRNAs in unperturbed MEFs and 136 miRNAs in unperturbed mESCs. These measurements showed that in these cells most miRNAs are long-lived, which confirmed the prevailing view of miRNA stability previously drawn from low-throughput analyses of a few miRNAs as well as broader analyses of cells perturbed with transcriptional inhibitors. The median half-life observed for miRNAs in dividing MEFs was 25 h—a value substantially greater than the 2.2 h median observed for mRNA half-lives in a similar cell type (dividing NIH3T3 cells) (T Eisen, D Bartel, unpublished). This >10-fold difference implies that levels of mRNAs can change more rapidly than those of miRNAs, making mRNAs much more adept at responding quickly to rapid environmental changes or other signaling cues, and relegating miRNAs to a more supporting role in lowering the half-lives of many mRNAs thereby helping mRNAs to achieve this more rapid response. The long half-lives generally observed for miRNAs are nonetheless suitable for changes over longer timeframes, such as those typically operating over the course of mammalian development, and they provide regulatory stability to the cell. Perhaps most importantly, they enable miRNAs to reach the high intracellular levels needed to impart consequential regulation, which are typically >1000 molecules per cell—levels much higher than the median mRNA level of ~17 molecules per cell (Schwanhäusser et al. 2011; Denzler et al. 2016).

Although most miRNAs were long-lived, our analysis identified a few with half-lives resembling those of intermediate-to long-lived mRNAs (1.7–10 h), including miR-503, a miRNA whose shorter half-life is proposed to facilitate its role in facilitating exit from cell-cycle arrest (Rissland et al. 2011). Among these relatively short-lived miRNAs, the mechanism of destabilization is known for only one, miR-7, a TDMD substrate. Indeed, in contact-inhibited MEFs, removal of the Cyrano lncRNA, which triggers miR-7 TDMD, increased the level of miR-7 by ~50-fold. Although the mechanism of destabilization of the other short-lived mRNAs is unknown, the observation that the miRNA nucleotides most important for target recognition also appeared to be most important for dictating half-life suggested a role for target pairing, as observed for miR-7 destabilization.

Combining metabolic labelling with approach-to-equilibrium kinetics enabled production rate constants to be determined alongside of decay rate constants, which provided a more complete understanding of miRNA dynamics than typically achieved using pulse–chase kinetics. These rate constants revealed that miRNAs can be produced at impressively rapid rates, both in proliferative as well as non-proliferative cells. The most abundant miRNA in dividing MEFs, miR-21a-5p, was produced at a rate of 114 ± 49 copies/cell per min. Even at a production rate of 24 copies/cell per min (two standard deviations below the calculated rate), the rate of miR-21a production would be more than 2-fold faster than that of the most rapidly produced mRNA in NIH3T3 fibroblasts and comparable to that of pre-rRNA production from a single pre-rRNA locus in HeLa cells (Schwanhäusser et al. 2011; Turowski and Tollervey 2015). When considering that miR-21 is transcribed from two alleles of a single locus and that RNA polymerase II (PolII) elongates with a rate constant of ~4.3 kb/min and has a footprint of ~40-50 nucleotides (Darzacq et al. 2007; Darst et al. 1991; Rice et al. 1993; Krebs et al. 2017), our results imply that at the calculated rate of production, the *Mir21* locus is coated in elongating PolII at approximately 50% of its maximum density. Such efficient PolII recruitment is presumably challenging and greater then 2-fold more efficient recruitment would be impossible, which helps explain why some highly expressed miRNAs are transcribed from multiple genes—the extreme being miR-430, which makes up 99% of the miRNA in the early zebrafish embryo and is transcribed from an array of >90 genes (Wei et al. 2012; Giraldez et al. 2005).

Our experiments also provided insight into previously inaccessible portions of the miRNA life cycle. Investigation of the dynamics of passenger-strand turnover, using the biexponential fit designed to distinguish between molecules acting as passenger strands and those loaded as guides into AGO, revealed that the combined processes of duplex loading and silencing-complex maturation, with slicing or expulsion of the passenger strand, can occur in < 1 min and typically within 10 min. These results indicate that the much longer lag time observed for the action of synthetic siRNA duplexes arises from other steps of that pathway, such as entry into cells and endosome release (Wittrup et al. 2015).

The biexponential model applied to assess passenger-strand dynamics assumed that one strand from every duplex succeeded in being loaded into AGO, with no degradation of miRNA duplexes prior to loading into AGO. However, in the context of miRNA over-expression, increasing AGO expression leads to increased miRNA abundance (Diederichs and Haber 2007), suggesting that in conditions of artificially increased miRNA levels surplus duplex exists and is degraded before either strand can be loaded as the guide. Although the prevalence of duplex degradation in the context of endogenous miRNA levels is unknown, the concentration of free AGO might be limiting for at least some miRNA duplexes in the cell. The observation that guide and passenger strands have such divergent half-lives implies that such duplexes, if they exist, degrade rapidly compared to the degradation rate of loaded guide strand, in which case degradation of unloaded duplex would not be expected to distort the guide-strand half-life values. With respect to passenger-strand values, if the unloaded duplexes degrade more rapidly than the time required for duplex loading and silencing-complex maturation, then the premature duplex degradation would reduce our passenger-strand half-life values but not substantially (unless a surprisingly large fraction of the duplex was degraded before loading). Conversely, if the unloaded duplexes degrade less rapidly than the time required for duplex loading and silencing-complex maturation, then the premature duplex degradation would increase our passenger-strand half-life values, and the values that we report would represent upper limits on the time required for duplex loading and silencing complex maturation.

Some studies have reported effects of certain terminal modifications on individual miRNAs (Jones et al. 2009; Katoh et al. 2009; Mansur et al. 2016), and in many systems, guide-RNA degradation observed upon loss of terminal 2′-O methylation is associated with tailing or trimming (Li et al. 2005; Ameres et al. 2010; Kamminga et al. 2010; Kamminga et al. 2012; Lim et al. 2015). However, the overall role of trimming and tailing in the degradation pathway of mature metazoan miRNAs had not been investigated. Here, we applied a model of isoform dynamics to extract rate parameters for mature miRNA production, mature miRNA conversion to trimmed and tailed species, as well as disappearance of those trimmed and tailed species. The broad spread of rate constants for each of these processes reflected the miRNA-specific nature of isoform dynamics, with different miRNAs differentially acquiring terminal modifications and then differentially responding to these modifications. Despite this variation, the addition of a single U both occurred at the fastest rate and was associated with the greatest degree of destabilization. Nonetheless, changes in isoform dynamics observed between MEF cell states poorly reflected the changes in miRNA half-lives, which implied substantial independence of the two pathways. Thus, our results suggested that trimmed and tailed species generally do not represent intermediates of the miRNA decay pathway, although these results do not exclude the possibility that for select miRNAs trimming or tailing might accelerate degradation. As these isoform analyses were carried out only on MEF datasets, we also cannot eliminate the possibility that changes in trimming and tailing might play a role more globally in altering stabilities between different cell types. Furthermore, as we could not fit the rate constants for isoform disappearance and back-conversion independently, it is also possible that more significant correlations between differences in miRNA stability and either of these parameters could have been obscured. However, the proposed independence of the two pathways is more broadly supported by the observations that loss of the non-canonical poly(A) polymerase TENT2 (GLD2/PAPD4) does not significantly change miRNA levels in the mouse hippocampus or in THP-1 cells (Burroughs et al. 2010; Mansur et al. 2016). Additionally, although TENT2 loss substantially diminishes tailing of miR-7, it does not impede Cyrano-mediated TDMD (Kleaveland et al. 2018).

Comparing rates of miRNA turnover in contact-inhibited MEFs, dividing MEFs, and mESCs revealed both specific changes in rates of turnover of individual miRNAs as well as general changes in miRNA turnover rates, as illustrated by the decreases in the median half-life from 34 h in contact-inhibited MEFs to 25 h in dividing MEFs and 6.3 h in mESCs. Both the miRNA-specific and the more general differences implied roles for miRNA-extrinsic factors in regulating half-life. With respect to miRNA-specific differences, the observation that miRNA nucleotides most important for target interaction were also most associated with shorter half-lives suggested a role for targets in this specification, although with the exception of Cryano-triggered TDMD of miR-7a-5p, further studies will be required to identify targets responsible for these differences. With respect to the general differences observed between the cell states and cell types, the differential activities of more broad-spectrum but as-yet-unidentified decay factors presumably mediate the differences. The observation that AGO2 often outlived the miRNA, especially in mESCs, indicated that most miRNAs can be degraded in a way that allows AGO to be recycled for use with another miRNA. Interestingly, the much faster dynamics exhibited by miRNAs in mESCs as compared to those in either MEF state presumably poises mESCs for differentiation and the more rapid changes in miRNA levels that this entails. The discovery of these miRNA-specific differences and the more general differences in miRNA dynamics observed between cell types will facilitate identification of the miRNA-extrinsic factors that mediate these differences. It also lays the foundation for further exploration of regulated miRNA turnover and how it interfaces with regulation of miRNA production to help drive or reinforce biological transitions.

## Materials and Methods

### Cell culture

MEFs were collected from E13.5 embryos of wild-type and Cyrano-knockout C57BL/6J mice (Kleaveland et al. 2018). Freshly harvested embryo bodies were minced in 1 mL 0.25% Trypsin-EDTA (Life Technologies) and incubated at 37°C for 30–45 min. Trypsin was then quenched by adding 4 mL MEF media (Dulbecco’s Modified Eagle Medium (DMEM, VWR) with 10% Fetal Bovine Serum (FBS, Takara) and 100 U/mL penicillin/streptavidin (pen/strep, Gibco)), and cells were dissociated by pipetting up and down. Suspended cells were transferred to a 10 cm dish, 6 mL of MEF media was added, and cells were allowed to grow to confluency before initiating passaging. MEFs were then immortalized with a retrovirus expressing the HPV type 16 E6/E7 genes and the selectable marker neomycin (Halbert et al. 1991). The retrovirus was produced by PA317 LXSN 16E6E7 cells grown in DMEM with high glucose (4 mM Glutamine), and purified from the cellular supernatant by centrifugation (3000 rpm for 15 min) followed by filtration through a Millex-HV 0.45 μm syringe filter. The retrovirus was then tittered to determine MOI and added to the MEFs at an MOI of 0.004 to obtain a polyclonal population with >99% single integrations. Immortalized cells were selected by continued passaging in the presence of the antibiotic G418 for three weeks. Following immortalization, MEF lines were grown at 37°C, 5% CO_2_ in DMEM and 10% FBS. MEFs were passaged every 2–3 d, using 0.25% Trypsin-EDTA for dissociation.

The v6.5 mESC line was cultured at 37°C, 5% CO_2_ on plates coated with 0.2% gelatin (Sigma, St. Louis, MO) in 2i media (1× DMEM/F12 (Gibco), 0.5× B27 supplement (Gibco), 0.5× N2 supplement (Gibco), 0.5× GlutaMAX (Thermo Fisher Scientific), 1× MEM NEAA (Gibco), 0.0025% BSA Fraction V (Gibco), 0.1 mM BME (Sigma), 100 U/mL U Pen/Strep, 1000 U/mL LIF (Millipore), 1μM PD0325901 (Stemgent), 3 μM CHIR99021 (Stemgent)). Cells were passaged and split 1:20 every 2 days. For passaging, cells were first washed with −CaCl_2_, −MgCl_2_ PBS (Thermo Fisher Scientific) and then dissociated with TrypLE Express (Thermo Fisher Scientific). Dissociated cells were resuspended in Serum/LIF media (1× DMEM KO (Gibco), 15% FBS, 1× MEM NEAA, 100 U/mL Pen/Strep, 1× GlutaMAX, 0.11 mM BME, and 1000 U/mL U LIF), pelleted by spinning at 1000g for 3 min, and resuspended in DMEM/F12 media for plating. HEK293FT cells (Thermo Fisher Scientific) were cultured at 37°C, 5% CO_2_ in DMEM and 10% FBS and passaged every 2–3 days, using 0.25% Trypsin-EDTA for dissociation.

Cells were counted using a Countess automated cell counter (Invitrogen). To determine doubling times for the different cell lines, cells were plated and then counted at intervals following plating. These data were then fit to a single-exponential function to extract doubling time, which was determined to be 22.48 h for dividing MEFs and 9.34 h for mESCs. The doubling time for contact-inhibited MEFs was assumed to be infinite.

### 5EU labeling and cell collection

For experiments examining contact-inhibited MEFs, cells were plated in 15 cm dishes and allowed to reach confluency. Cells were then left confluent for 4 days, with media changes every 2–3 days. On the fifth day of confluency, a media change was performed, which was timed such that all plates that were to be collected ≤ 24 h after EU addition were last fed 24 h prior to collection. The following day, 5EU (Jena Bioscience) was added to the culture media to a final concentration of 400 μM, and cells were collected 0, 1, 2, 4, 8, 24, 72, and 168 h later. For time intervals longer than 24 h, the media and 5EU were refreshed every 24 h, timed such that the last feeding was 24 h prior to collection. A total of 14 dishes were plated for each experiment, with one plate each for the 0, 24, 72, and 168 h time intervals, 2 plates each for the 4 and 8 h time intervals, and 3 plates each for the 1 and 2 h time intervals.

For the experiments examining dividing MEFs, cells were plated 24 h in advance of collection at a density of either 1.25 million (replicate 1) or 1 million (replicate 2) cells per 15 cm plate. 5EU was added to a final concentration of 400 μM, and cells were collected at 0, 1, 2, 4, 8, 24, and 72 h. During the 72 h time interval, cells were split and re-plated daily with fresh media and 5EU. The same number of plates were used for each time interval as were used for the contact-inhibited MEFs.

For the mESC experiments, cells were plated 48 h in advance of collection at a density of 200,000 cells per 15 cm dish. The next day, media was changed for the 0, 1, 2, 4, and 8 h time intervals 24 h prior to planned collection. Media was also changed 24 h in advance for the 16 and 24 h time intervals, and 5EU was added to a final concentration of 125 μM. The following day, 5EU was added at the same concentration for the remaining time intervals, and cells were collected.

For all experiments, cell collection proceeded by adding TRI Reagent (Life Technologies, 3 mL per 15 cm dish), scraping the cells off the plate, and then transferring this mixture to a 15 mL conical tube which was then snap frozen and stored at −80°C. For the dividing MEFs, cells were washed with PBS prior to the addition of TRI Reagent. For the mESCs, cells were similarly washed with −CaCl_2_, −MgCl_2_ PBS (Gibco).

### RNA extraction and small RNA enrichment

Samples were thawed and then phase-separated with the addition of chloroform (J.T. Baker Analytical) at a ratio of 250 μL per 1 mL TRI Reagent. RNA was then precipitated with isopropanol, and pellets were washed twice with 70% ethanol prior to resuspension in water. Following RNA extraction, the quantitative standards were added to a level of 1 fmole/20 ng of total RNA (contact-inhibited MEFs, replicate 1), 1 fmole/10 ng of total RNA (dividing MEFs replicate 1), or 1 fmole/1 ng of total RNA (contact-inhibited and dividing MEFs, replicate 2, mESCs). For the MEF samples, small RNAs were then enriched from the total RNA sample with the miRvana miRNA isolation kit (Thermo Fisher Scientific). Following small-RNA enrichment, samples were precipitated and resuspended in water.

### Biotinylation and pulldown

Biotin was attached to metabolically labeled RNAs in a 10–20 μl reaction with 4 mM biotin disulfide azide (Click Chemistry Tools), 5 mM CuSO_4_ (Sigma Aldrich), 5 mM THPTA (Click Chemistry Tools), 20 mM Sodium L-Ascorbate (Sigma Aldrich) and 50 mM HEPES, pH 7.5. After incubating for 1 h at room temperature with protection from light, the reaction was then quenched with 5 mM EDTA, and RNA was extracted using phenol chloroform (Sigma Aldrich) and precipitated. For the pulldown, 100 μL MyOne Streptavidin C1 bead slurry (Life technologies) was used for every 25 μg RNA in the click reaction. The beads were washed twice with B&W buffer (10 mM Tris, pH 7.5, 1 mM EDTA, 2 M NaCl, and 0.01% Tween 20), twice with solution A (0.1 M NaOH, 0.05 M NaCl, 0.01% Tween 20), twice with solution B (0.1M NaCl, 0.01% Tween 20), and twice with water. Beads were then blocked for 30 min at room temperature on an end-over-end rotator with 0.5 μg/μL of yeast total RNA diluted in high-salt wash buffer (HSWB) (10 mM Tris, pH 7.5, 1 mM EDTA, 100 mM NaCl, and 0.01% Tween 20). Blocked beads were then washed three times with HSWB. The RNA pellet was dissolved in HSWB (100 μL per 100 μL of beads used), and this solution was used to resuspend the blocked and washed beads. After incubation on an end-over-end rotator for 30 min at room temperature, beads were washed twice with 50°C water, and then twice with 50°C 10X HSWB. During the final wash, beads were transferred to a new tube, the final wash was removed, Tris(2-carboxyethyl)phosphine hydrochloride (TCEP, Sigma Aldrich) was added (200 μL per 100 μL beads), and the beads were incubated on an end-over-end rotator at 50°C. After 20 min, the TCEP eluate was moved to a new tube, the beads were washed once with 150 μL of water, the TCEP eluate and wash were pooled, NaCl was added to a concentration of 0.3M, and RNA was precipitated with 2.5 volumes of 100% ethanol.

Pulldown selectivity and efficiency was assessed by examining the behavior of the quantitative radiolabeled standards. Ten percent of the input, flow-through, and elution fractions from the pulldown were run out on a 15% polyacrylamide urea gel, and recovery of the standards was quantified by phosphorimaging (Typhoon FLA 7000, GE Healthcare). Enrichment of EU-containing standards was also assessed for each timepoint from the sequencing data.

### Generating EU-containing quantitative standards

DNA oligos used for in vitro transcription (standard and T7 template strands in Supplemental Table S3) were annealed in a 100 μl reaction containing 40 μM of each oligo and 0.15 M NaCl, which was heated to 95°C for 5 min and then cooled to room temperature. Annealed duplexes were diluted to 1 μM in 0.1M NaCl and transcription was then carried out with the T7 MEGAshortscript kit (Thermofisher Scientific). To label standards with 5EU, 5EUTP (Jena Bioscience) was diluted to 75 mM in water and used in place of the UTP stock solution. After incubation at 37°C for 4 h, transcription reactions were phenol–chloroform extracted, and the desired transcript was purified on a 10% polyacrylamide urea gel and resuspended in water at a concentration of 5 μM.

Five pmol of each standard was dephosphorylated in a 100 μL reaction of 1× CutSmart Buffer (NEB) with 5 units of calf intestinal alkaline phosphatase (CIP, NEB) for 30 min at 37°C, phenol–chloroform extracted to quench the CIP reaction, ethanol precipitated to reduce the volume to 7 μL, and then labeled, according to the manufacturer’s protocol, with T4 Polynucleotide Kinase (PNK, NEB) and 1μL of 25 μM γ^32^P-ATP (Perkin Elmer) for at least 1 h at 37°C. Labeled samples were de-salted with a P30 column (Bio-Rad laboratories) and then purified on a 10% polyacrylamide urea gel.

### Small-RNA sequencing

Samples for library preparation were assembled by mixing either 2–5 *μ*g of total RNA (input libraries) or the entire eluate from 5EU pulldown (EU libraries) with size-selection markers and quantitative sequencing standards. Size-selection markers were 18 and 32 nt 5'-end-labeled RNAs (Supplemental Table S3). Sequencing standards (Supplemental Table S3) were added at a level of either 0.1 fmol per 1 *μ*g total RNA (input libraries) or 2.5 amol per sample (EU libraries). Both 3' and 5' ligations were carried out with degenerate adaptors (each containing 4 random nucleotides) in the presence of 10% Polyethylene Glycol (PEG 8000, NEB) and 0.5 *μ*L of Superasin (Thermo Fisher Scientific) with either T4 RNA-ligase 2, truncated K227Q (NEB) or T4 RNA-ligase 1 (NEB), respectively. Reverse transcription was with SuperScript III (NEB), and subsequent PCR amplification of the cDNA was with Phusion (NEB). A step-by-step protocol for preparing small-RNA sequencing libraries is available at http://bartellab.wi.mit.edu/protocols.html. Samples were sequenced on an Illumina HiSeq platform with 40 nt single-end reads.

Sequencing reads were trimmed at the 5' and 3' ends using fastx_trimmer (FastX Toolkit; http://hannonlab.cshl.edu/fastx_toolkit/) and cutadapt (Martin 2011), respectively. Trimmed reads were subsequently filtered for greater than 99.9% accuracy for all bases using fastq_quality_filter (FastX Toolkit) with the parameters ‘-q 30 −p 100’. To call miRNA reads, the first 19 nucleotides of filtered and trimmed reads were string-matched to a dictionary of miRNA sequences. The miRNA dictionary was curated by filtering miRbase_v21 miRNA annotations for all conserved or confidently annotated small RNA species (as annotated by TargetScanMouse, release 7.2) as well as their passenger-strand partners (Supplemental Tables S1,2). Although most miRNAs have a unique initial 19 nucleotides, a few species cannot be called unambiguously using only the first 19 nucleotides. These species were collapsed into a single dictionary entry (whose sequence was chosen randomly to be one of the collapsed species’ sequence) and listed under a merged name (for example, mmu-miR-199a-3p and mmu-miR-199b-3p become mmu-miR-199ab-3p). Assigned reads were normalized within each time course using either the EU-containing quantitative standards (EU data) or the sequencing standards (input data). Each miRNA in the normalized EU dataset was then further normalized to its abundances across the normalized input dataset to account for any potential non-steady-state behavior. With the exception of the passenger and guide comparisons, all analyses were carried out with only the guide strands; guide strands were annotated based on abundance in the input libraries.

For trimming and tailing analyses, suffixes to the 19 nucleotide prefixes were enumerated for each miRNA. All suffixes representing more than 20% of the reads for a miRNA in the 1 h EU time interval were designated initial products of biogenesis (Supplemental Table S1), and the trimmed and tailed suffixes were annotated with respect to this initial isoform. Only guide RNAs with one initial isoform were used when modeling isoform dynamics.

### RNA sequencing

RNA-seq samples were prepared using the NEXTflex Rapid Directional mRNA-seq Kit (Bioo Scientific). Briefly, mRNAs were enriched from 10 μg total RNA with NEXTflex Poly(A) Beads (Bioo Scientific). These species were then fragmented, synthesized into cDNA, ligated to adaptors, and PCR amplified as per the manufacturer’s protocol. Samples were then sequenced on an Illumina HiSeq platform with 40 nt single-end reads.

Reads from MEFs and mESCs were aligned to the mouse genome (mm10) with Star v. 2.4 with the parameters “--alignIntronMax 1 --runThreadN 30 --outFilterMultimapNmax 1 -- outFilterMismatchNoverLmax 0.04 --outFilterIntronMotifs RemoveNoncanonicalUnannotated -- outSJfilterReads Unique”. These reads were then assigned to genes based on annotations from Ensembl (Mus_musculus.GRCm38.94.gtf downloaded September 5, 2018) with htseq-count v0.9.1(Anders et al. 2015) with the parameters ‘-m union −s reverse’. Targets present in the different cell lines were identified and classified based on TargetScanMouse v7.2 annotations. Raw RNA-seq reads from mouse 3T9 fibroblasts (Ghini et al. 2018) were downloaded from the GEO (GSE104650) and processed as described above, except the parameters ‘-m union −s no’ were used for htseq-count, which accommodated the non-stranded 3T9 dataset.

### Northern blotting

After resolving 5–10 μg of total RNA on a 20% polyacrylamide urea gel, RNA was transferred to a Hybond-NX membrane (GE Healthcare) with a semi-dry transfer apparatus (Bio-Rad) and crosslinked to the membrane by incubation with EDC (*N*-(3-dimethylaminopropyl)-*N*′-ethylcarbodiimide; Thermo Scientific) diluted in 1-methylimidazole for 1–2 h at 60°C. Blots were probed with either DNA or LNA oligonucleotide probes (Supplemental Table S3). Results were quantified using ImageQuant TL (v8.1.0.0). A step-by-step protocol can be found at http://bartellab.wi.mit.edu/protocols.html.

### Pulse-labeling with heavy amino acids and AGO isolation

SILAC media were prepared essentially as described (Ong and Mann 2006). DMEF base media was made by mixing DMEM lacking Lysine and Arginine (Thermo Fischer Scientific) with dialyzed FBS (Life Technologies). Base media for mESCs was made with DMEM/F-12 lacking Lysine and Arginine (Life Technologies) plus all additional components required for mESC culture as listed above. These base media were then supplemented with 84 mg/mL ^13^C_6_^15^N_4_ L-arginine plus 146 mg/mL ^13^C_6_^15^N_2_ L-lysine or the corresponding non-labeled amino acids (all from Sigma) to generate either heavy or light media, respectively. MEFs and mESCs were grown in light SILAC medium for at least 5 passages prior to pulse-labeling. Contact-inhibited MEFs were seeded onto 10 cm dishes and allowed to reach a contact-inhibited state 5 days prior to the start of the pulse labeling, whereas dividing MEFs and mESCs were plated at a density of 225,000 cells per 10 cm dish or 200,000 cells per 15 cm dish, respectively, 48 h prior to the planned time of collection. At time zero, the light media was removed, and cells were washed three times with pre-warmed PBS before addition of transfer into heavy media. Cells were collected 1.5, 4.5, 14, and 24 h after growth in heavy media and washed twice with ice-cold PBS, scraped off the plate, spun down, and snap frozen.

Frozen pellets were resuspended in 500 μL of NET buffer (50 mM Tris, pH 7.5, 150 mM NaCl, 5 mM EDTA, 0.5% NP−40, 10% glycerol, 1 mM Sodium Orthovanadate, 0.5 mM DTT; supplemented with cOmplete EDTA-free Protease Inhibitor Cocktail (Roche)) and incubated at 4°C for 20 min. The lysate was then passed through a 23G syringe seven times before centrifugation at 15,000g for 20 min. AGO proteins were enriched based on their affinity to a peptide derived from TNRC6 (T6B) as described (Hauptmann et al. 2015). Briefly, Protein G beads (Life Technologies) were washed once with PBST (PBS, 0.02% Tween) before being incubated with anti-FLAG antibody (Sigma Aldrich) at room temperature with end-over-end rotation for 10 min. The beads were then washed twice with PBST before incubation with FLAG-tagged T6B peptide at 4°C with rotation for 2 h. Peptide-coupled beads were washed three times with PBS, and then resuspended in cell lysate and rotated at 4°C for 3 h. Beads were then washed four times with NET buffer, once with PBS, and then resuspended in NuPAGE 2× LDS Sample buffer (ThermoFischer Scientific).

### Mass spectrometry and AGO2 half-life estimation

AGO-enriched samples were incubated at 95°C 5 min and then resolved on a 4-12% Bis-Tris NuPAGE gel (Thermo Fischer Scientific). Gels were stained with Imperial Protein Stain (Thermo Fischer Scientific), and a band migrating near 100 kDa was excised, cut into ~2mm squares, and washed overnight in 50% methanol. The following day the gel pieces were washed once more with a solution of 47.5% methanol, 5% acetic acid for 2 h before being dehydrated with acetonitrile and dried in a speed-vac. Samples were then incubated in 100 mM ammonium bicarbonate with 10 mM dithiothreitol for 30 min to reduce disulfide bonds, before subsequent incubation in a solution of 100 mM ammonium bicarbonate and 100 mM iodoacetmide for Cysteine alkylation. Samples were then sequentially washed with acetonitrile, 100 mM ammonium bicarbonate and acetonitrile and dried in a speed-vac. Proteins were subsequently digested in a solution of 50 mM ammonium bicarbonate with 20 ng/μL trypsin; following addition of the trypsin, samples were first incubated on ice for 10 minutes before digestion overnight at 37°C with gentle shaking. The digested peptides were extracted by serial 10-minute incubations at 37°C with shaking in 50 mM ammonium bicarbonate, a solution of 47.5% acetonitrile, 5% water, and 5% formic acid, and the acetonitrile solution once again. Following each incubation step, the supernatant was removed and set aside. These supernatants were subsequently pooled, the organic solvent was removed, and the sample volumes were reduced to 15 μl using a speed vac.

Samples were next analyzed by reversed phase high performance liquid chromatography (HPLC) using a Thermo EASY-nLC 1200 HPLC equipped with a self-packed Aeris 1.7 μm C18 analytical column (0.075 mm by 14 cm, Phenomenex). Peptides were eluted using standard reverse-phase gradients, and effluent was analyzed using a Thermo Q Exactive HF-X Hybrid Quadrupole-Orbitrap mass spectrometer (nanospray configuration) operated in a data dependent manner. The resulting fragmentation spectra were correlated against the known database using PEAKS Studio X (Bioinformatic Solutions).

Heavy-to-light ratios were computed for all AGO2 peptides that had peak areas greater than 10,000 for both heavy and light peptides all timepoints. These ratios were used to determine the disappearance rate constant of AGO2 by fitting the data to the following model: *ln*(*r* + 1) = *k*_*d*_(*t* − *t*_0_) where *r* is the ratio of the peak areas for the heavy or light isotope of each individual peptide, *k*_*d*_ is the disappearance rate constant, *t* is the time, and *t*_0_ is the time offset due to the lag in heavy isotope incorporation following the media change (Schwanhäusser et al. 2011). The parameters *k*_*d*_ and *x*_0_ were fit by using the optim function in R (method="L-BFGS-B", R version 3.2.5) to minimize the sum of the squared residuals of this model to the data. All fit parameters were bounded to be > 10^−8^. As *k*_*d*_ is the sum of the rate of dilution due to cell division and the rate of protein degradation, the AGO2 degradation rate constant (*k*_*deg*_) was determined by subtracting the rate of dilution due to cell division (*ln*(2)/*t*_*cc*_, where *t*_*cc*_ is the duration of a single cell cycle) from the fit value for *k*_*d*_. Values of *t*_*cc*_ for each cell-type were determined empirically. AGO2 half-life was then calculated from *k*_*deg*_ as *t*_1/2_ = *log*(2)/*k*_*deg*_.

For comparison to AGO2 half-lives, a single representative half-life was calculated for the combination of all miRNAs in a given cell state or type. To generate the appropriate combination of all guide strands, we had to account for the slight U bias attributable to the pulldown procedure. This correction was accomplished with a U-bias scaling factor determined from the ratio of the theoretical steady-state value (calculated from abundance in the input sample) to the fit steady-state value for each miRNA. The alpha parameters for each guide strand were scaled by this factor, and then, in combination with the beta parameters, were used to determine the theoretical ratios of 5EU-labeled miRNAs to unlabeled miRNAs over time, which represented the pool of miRNAs synthesized in the time following 5EU addition and prior to 5EU addition, respectively. These ratios were then fit in a manner identical to the SILAC heavy-to-light ratios to extract the representative miRNA half-life.

### Fitting models to data

Data were filtered for expression, requiring at least 60 reads per million total reads aligned to miRNAs for every timepoint. Data passing this threshold were normalized to the average of the three EU-containing quantitative standards. The data were additionally normalized by miRNA levels observed in the input libraries for each time interval, which accounted for fluctuations in steady-state levels of miRNAs over the course of the experiment. When combining data from replicates, data were batch corrected to account for differences in the preparation of the quantitative standards used for different replicates. Both these combined datasets as well as data from individual replicates were then fit to the single-exponential models. Pairs of guide and passenger strands were selected from the read-filtered dataset based on fulfillment of both the requirement that the steady-state level of the guide strand be at least 5-fold greater than that of the star strand, as well as the requirement that both strands map to a unique genomic locus. For all analyses, half-lives were calculated as *ln*(2) divided by the degradation rate constant.

When fitting these data, the optim function in R (method="L-BFGS-B", R version 3.2.5) was used to minimize the sum of the squared residuals of the log-transformed model to the log-transformed data. For each dataset, fitting was carried out globally, minimizing the sum of the residuals for all miRNAs with reads passing the expression threshhold, which enabled extraction of a single value for the time-offset parameter (*t*_0_) for the entire dataset. For the single-exponential model, the values for the rate constants of production (*α*) and degradation (*β*) for each guide RNA were fit in addition to the global *t*_0_ value. The fit values for these rate constants were bounded to be > 10^−8^ (units of reads per hour or h^−1^, respectively), and the fit value for the time offset parameter was required to be between 0.01 and 1 h. of degradation For the two-exponential model, the values for one rate constant of production (*α*), four rate constants of degradation(*β*_*n*_), and one fractional split (*f*) were extracted for each guide–passenger pair, in addition to the global *t*_0_ value. To allow for the paired optimization of fits to guide and passenger strand data, the U-bias effects were accounted for by scaling the production rates for all passenger and guide strands by the U-bias scaling factor. As a result of this scaling factor, the fit production rates reported for guide and passenger strands from a given pair are not identical. The bounds for the production and degradation rate constants as well as for *t*_0_ were the same as for the single-exponential models, and the value for the fractional split was bounded to be between 10^−8^ and 1. For all models, the rate of cell division (*g*) was manually set based on experimental determination. To ensure that the true minima was reached by the optim function, optimization was run iteratively 100 times for each dataset using the parameter values reached by the previous round of optimization as the starting values for the next round.

For isoform analyses, miRNA behavior across time was decomposed into time courses for individual isoforms. Due to sequencing depth and the low abundance of miRNA isoforms, for most miRNAs only the initial biogenesis products and their +1 (A), +1 (U), and −1 isoforms had substantial signal across the time course. All species with greater than one raw read for these isoforms at each time interval were fit with the isoform dynamics model. Replicates were combined with batch-correction (described above), and again, fitting was carried out to these combined time courses by using the optim function in R (method="L-BFGS-B", bounds set to 0.000001 and 1000000 for all parameters) to minimize the sum of the squared residuals. All isoforms for a given miRNA were fit simultaneously with a system of differential equations that was solved by numerical integration with the lsoda function. Fitting was carried out in log-space, and optim was iteratively run 10 times to ensure true minimization.

### Family analyses

All guide strands were assigned to families based on seed region sequence. Comparisons were then made between all pairs of family members within each family. Guide strands with capped half-lives (either > 72 h in dividing MEFs, or > 24 h in mESCs) were not considered in these analyses due to uncertainty of their true half-life values. The metrics of percent difference and hamming distance were used to measure half-life variation and sequence similarity, respectively. Random comparisons used equal numbers of non-family pairs, selected randomly, without replacement.

### Motif and feature searches

To search for motifs associated with rapid turnover, miRNAs were either ranked according to half-life or classified into fast- and slow-turnover subsets. To search for position-specific motifs, a list of miRNA sequences ranked by half-life was supplied to kpLogo (Wu and Bartel 2017). To query differences in more general nucleotide content, miRNAs were classified as fast or slow turnover. The fast- and slow-turnover subsets were delineated as species that had the upper or lower bounds of their 95% confidence intervals below or above certain thresholds (specified in the figure legend where applicable). Such classification ensured that the subsets compared were truly differentially turned over. Different features of these fast and slow turnover subsets were also compared to the analogous features for equal size subsets of all of the miRNAs examined in this study.

### Quantification and statistical analyses

Graphs were generated and statistical tests were performed using R (v3.2.5). Statistical parameters including the test used, significance ascertained (*p* value), and number of points compared (*n*) are reported in either the figures or their legends. For EU time courses, replicates refer to biological replicates carried out on different days. Standard deviations and confidence intervals on the fit parameters were determined by computing the covariance matrix for these parameters from the variance of the residuals and an approximation of the Hessian of the error function. Statistical tests were selected based on the comparison being made. No statistical tests were performed to pre-determine sample size. A two-sided Z-test was applied to identify miRNAs that exhibited significantly higher rates of production than expected for their abundance (significance required |Z-score| > 2). Based on the nature of the data, either an unpaired two-sample, two-tailed t-test or a Mann-Whitney U test was used for comparisons of the distributions of guide and passenger half-lives, half-life percent differences, and isoform rate constants. Significance of correlations between half-life percent difference and sequence similarity for family members was assessed using a correlation test (cor.test function, method = ‘pearson’, R v3.2.5).

### Data Access

All raw and processed sequencing data generated in this study have been submitted to the NCBI Gene Expression Omnibus (GEO; http://www.ncbi.nlm.nih.gov/geo/) under accession number GSEXXXXXX.

## Acknowledgements

We thank members of the Bartel lab for helpful discussions; T Eisen, J Kwasnieski, S McGeary, and K Lin for computational advice; B Kleaveland for sharing of reagents and samples, and the Whitehead Institute Genome Technology Core for sequencing. This work was supported by N.I.H. grant GM118135 (D.P.B.) and an N.S.F. predoctoral fellowship (E.R.K.). D.P.B. is an investigator of the Howard Hughes Medical Institute.

---

**Supplemental Figure S1.**
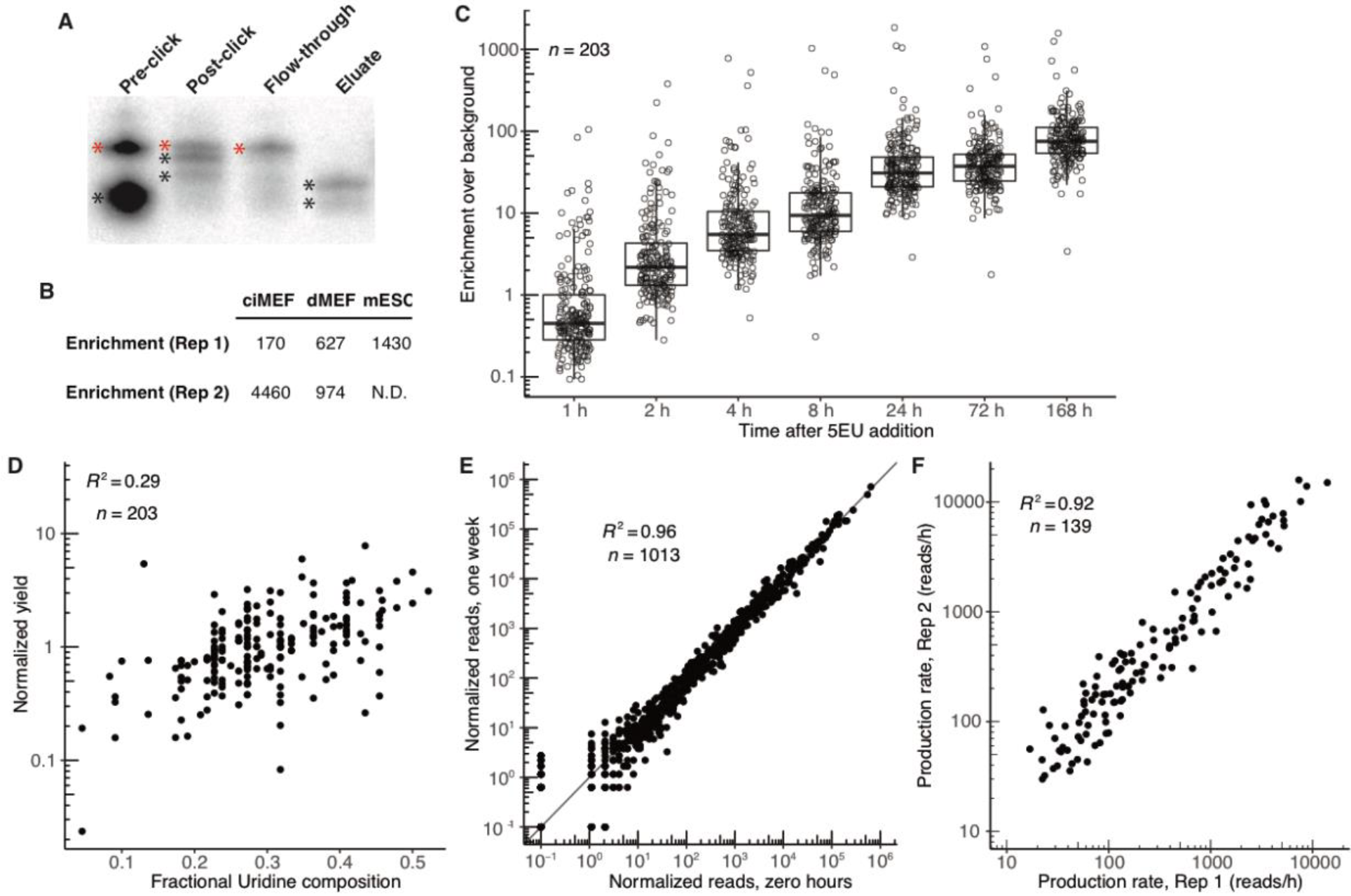
Labeling with 5EU allows efficient and selective pulldown of small RNAs with minimal background or bias. (A) Selective enrichment of 5EU-containing standards. Shown is a phosphorimager scan of a denaturing acrylamide gel that resolved the three radiolabeled 22–25-nt small RNAs that each contained a single 5EU (black asterisks) and an unlabeled 30-nt RNA (red asterisks). The pre-click lane shows the ratio of these species prior to biotinylation, and the post-click lane shows that the addition of the biotin group slowed mobility of the 5EU-containing standards. The other lanes show that the flow-through was depleted in 5EU-containing standards, whereas the eluate was strongly enriched in 5EU-containing standards, which had regained their faster mobilities because the biotin group was cleaved from the RNA during elution. (B) Selective enrichment of 5EU-containing standards, as indicated by sequencing. Reported are median fold-enrichment values of 5EU-containing standards over a standard that lacked 5EU, as determined by small-RNA sequencing of samples from contact inhibited MEFs (ciMEFs), dividing MEFs (dMEFs), and mESCs. N.D. not done. (C) Enrichment of each individual miRNA (including each passenger strand passing the expression threshold) over its background level across time in contact-inhibited MEFs. Background levels were determined by sequencing the RNA eluted after performing the pulldown procedure on RNA from cells that had not been metabolically labeled. (D) Relationship between pulldown yield and the number of U nucleotides in the miRNA. For each miRNA (including passenger strands) in the contact-inhibited MEF dataset, median-centered normalized yield is plotted as a function of fractional U composition. Pulldown yields were normalized by dividing the reads from the 1-week EU eluate sample by the reads from the 1-week input sample and then median centered; this normalized yield indicates the extent to which each miRNA was enriched or depleted in the 5EU dataset compared to expectation. (E) A comparison of miRNA reads in the one-week input sample to those in the unlabeled input sample (reads for all confidently-annotated guide and passenger strands are shown). Reads for both samples, which were from contact-inhibited MEFs, were normalized to the internal standards. (F) Correspondence of production rates of guide strands from the two contact-inhibited MEF replicates.

**Supplemental Figure S2.**
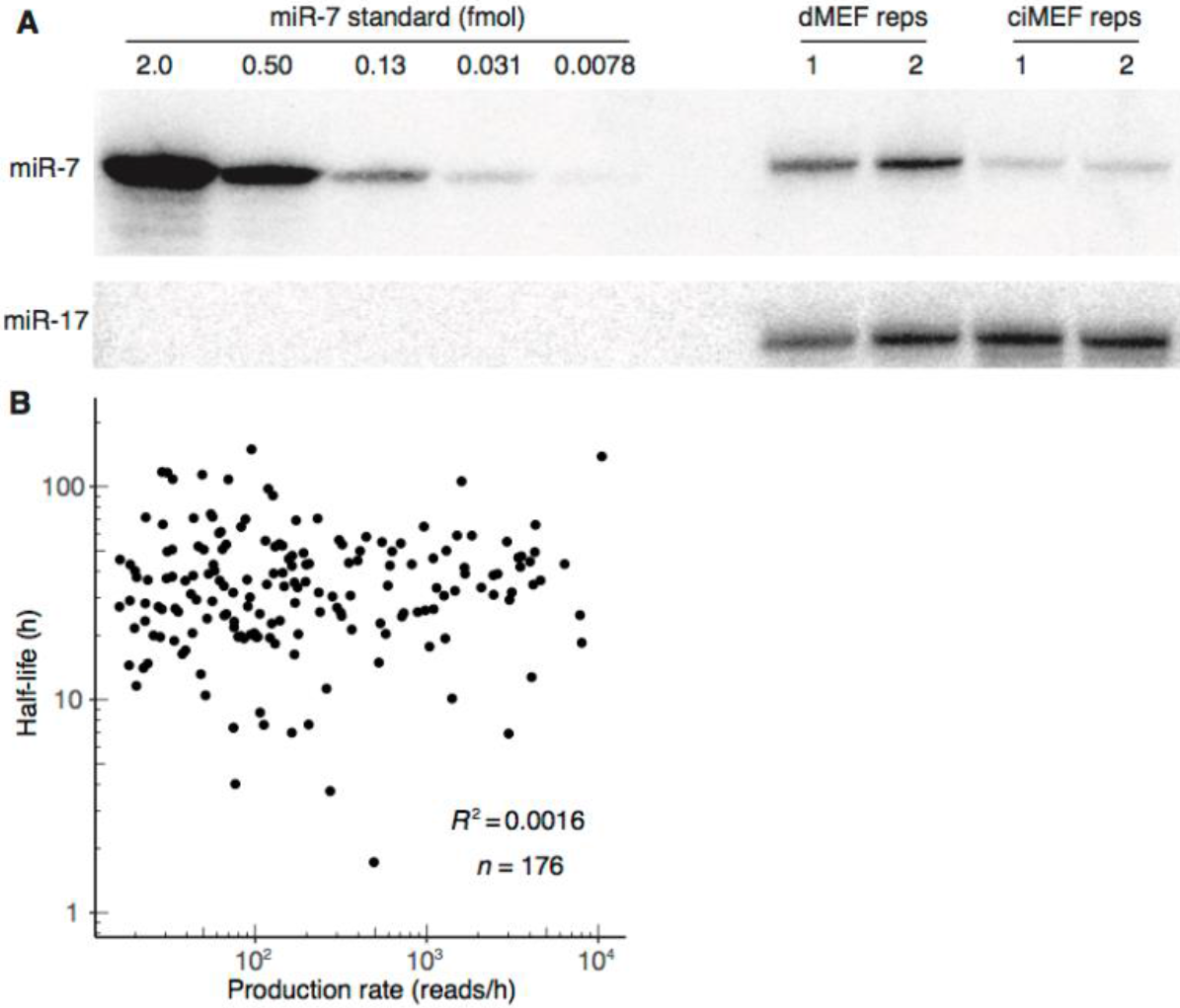
Dynamics of guide miRNAs in contact-inhibited MEFs. (A) Quantitative northern blot used to determine the absolute number of miR-7 molecules in contact-inhibited and dividing MEFs (ciMEF and dMEF, respectively). 10 *μ*g of total RNA, which corresponded to ~5 million cells, was loaded for each replicate (reps) for the dividing MEFs, and 20 *μ*g of total RNA, which corresponded to ~ 10 million cells, was loaded for each replicate for the contact-inhibited MEFs. Standards of the specified molar amounts were loaded for miR-7; miR-17 served as a loading control. (B) Relationship between half-life and production rate for guide strands in contact-inhibited MEFs.

**Supplemental Figure S3.**
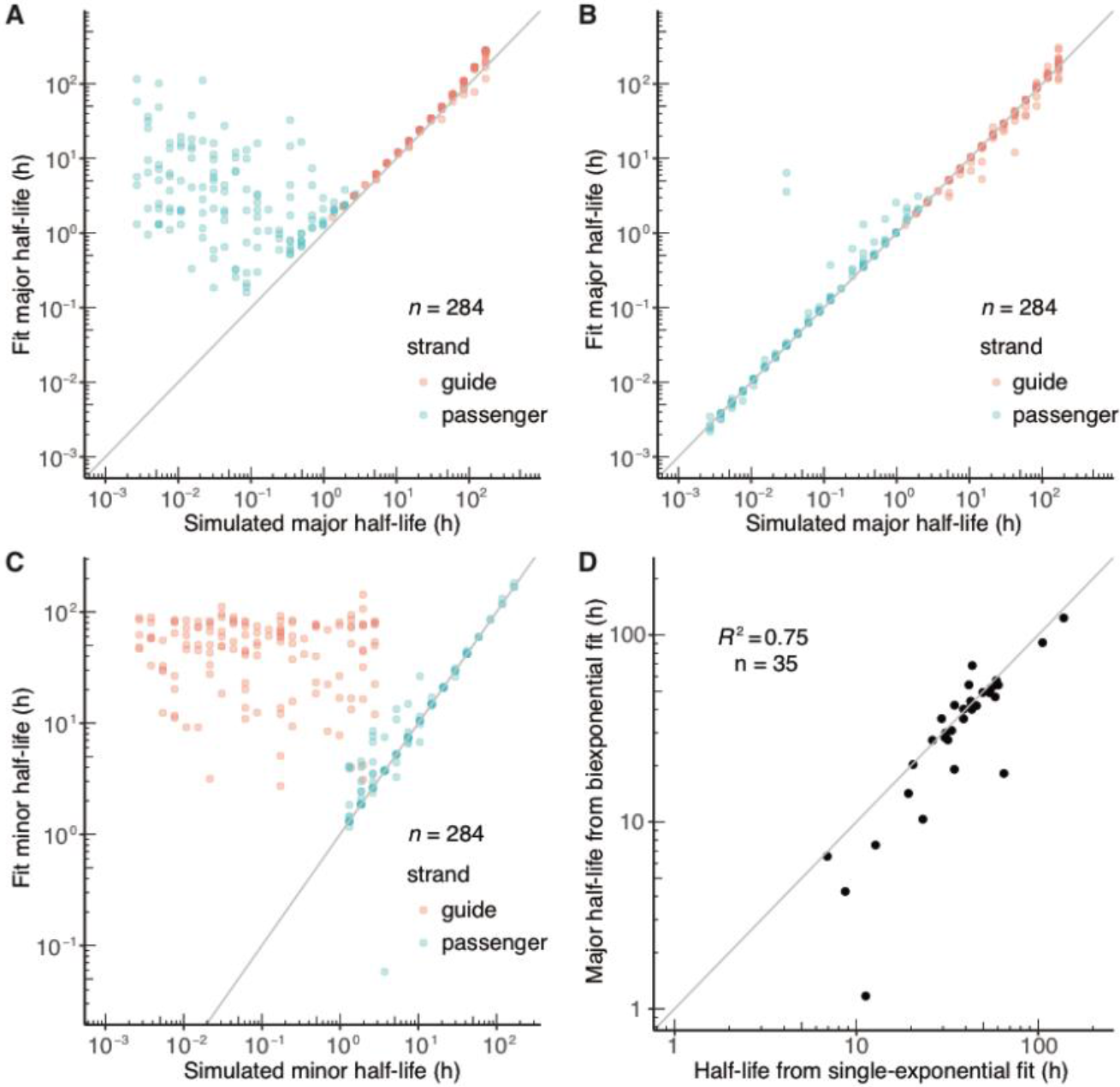
Fitting passenger and guide pairs to a biexponential model. (A–C) Assessing the ability of the single-exponential fits (A) and biexponential fits (B–C) to recapitulate simulated data for both guide and passenger strands. Bi-phasic behavior was simulated with a system in which each species was divided between two behavioral populations with a specified fractional distribution (Fig. 1B, eqn. 2). Both populations of each species were set to have the same production rate, but each had an individual rate of degradation (with the “major” and “minor” rates describing the turnover of the majority or the minority of the species, respectively) (Fig. 1B, eqn. 2). A wide array of different input parameters was fed into this model to generate many different bi-phasic behaviors. Initial simulations revealed the range of half-lives that the models could accurately fit, and these bounds were applied in the simulations shown here. Major half-lives from either single-exponential fits (A) or biexponential fits (B) were compared to the major half-lives used to generate the simulated data. Additionally, minor half-lives fit using the biexponential model were compared to the minor half-lives used to generate the simulated data (C). These simulations illustrated the utility of fitting to the biexponential model when determining the half-lives of the both the major and the minor passenger-strand populations. They also showed that fitting to either model was sufficient to determine half-life of the major guide-strand population (provided that the half-life did not exceed the longest time interval) and that fitting to neither model was reliable for determining the half-life of the minor guide-strand population. (D) Correspondence between half-lives determined by biexponential fits and half-lives determined by single-exponential fits for guide strands in contact-inhibited MEFs.

**Supplemental Figure S4.**
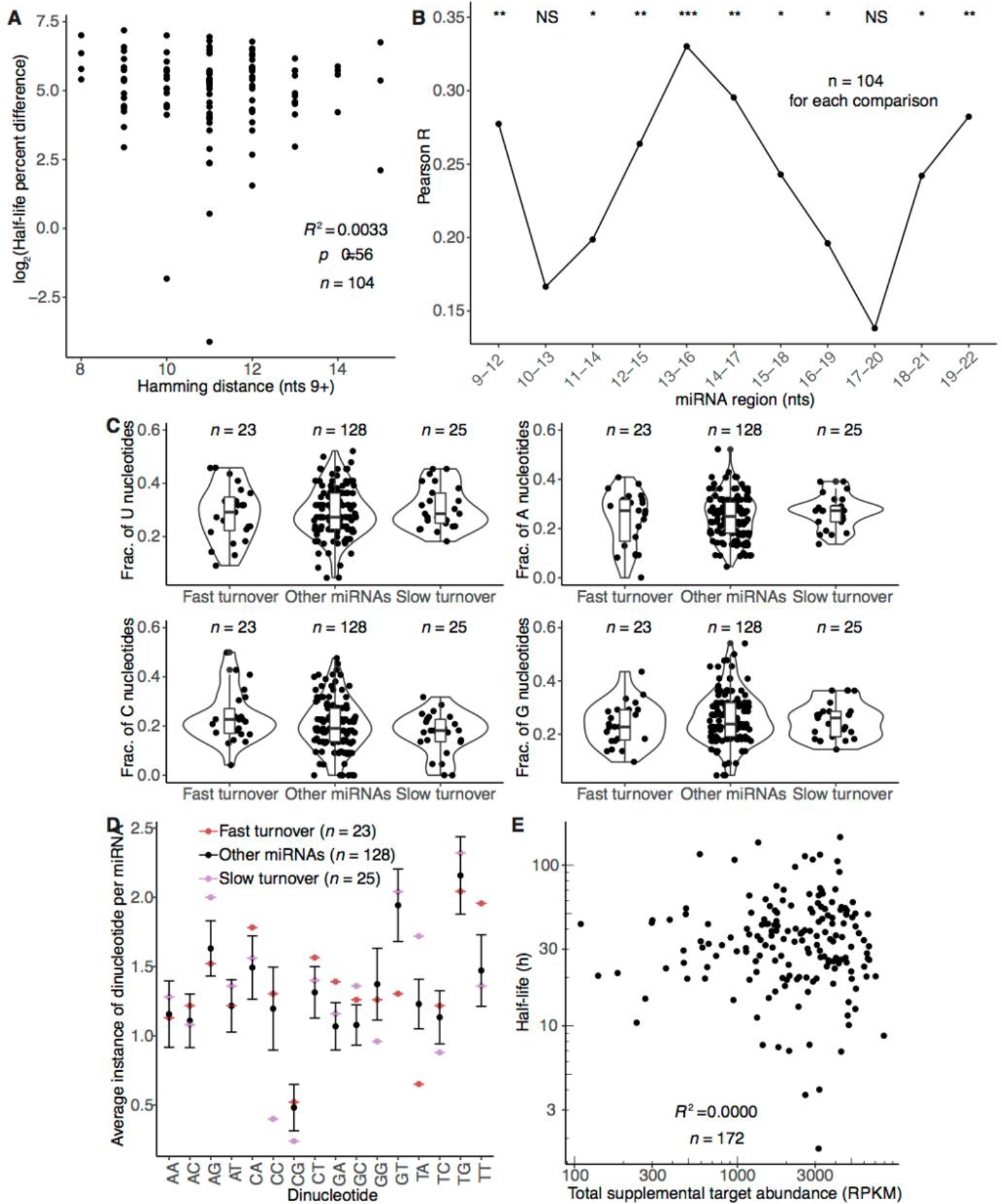
Search for determinants of turnover for guide strands in contact-inhibited MEFs. (A) Relationship between half-life difference and Hamming distance for random pairs of non-family members. Only nucleotides downstream of the seed region were considered for the Hamming-distance measurement. Significance of the correlation was determined by a t-test. Twenty-five equally sized cohorts of randomly paired guides were analyzed in the same way, with the plot shown here being for the cohort with the median *R*^2^ value. (B) Relationship between half-life percent difference and Hamming distance for 4-nt segments of family-member pairs. Plotted for each 4-nt segment downstream of the seed region is the Pearson *R* describing the correlation between these two metrics. Significance of each correlation was determined by a t-test. (C) Nucleotide composition for fast- and slow-turnover species as well as all other miRNAs. Fast-turnover species had half-life 95% confidence intervals that fell below 35 h; slow-turnover species had half-life 95% confidence intervals that fell above 25 h; other miRNAs fit into neither of these categories. (D) Average dinucleotide composition of fast-turnover miRNAs (red), slow-turnover miRNAs (purple), and other miRNAs (black). MicroRNAs were assigned to the categories as in (C). To allow comparison between equal-sized groups of miRNAs, the ‘other’ miRNAs were subset into random sets that were of equal size to the fast-turnover set. The dinucleotide composition was determined for 50 of these random cohorts, and the mean composition for all cohorts is shown (error bars, standard deviation). (E) The relationship between half-life and summed RPKM of all predicted targets with supplemental pairing to the miRNA. Predicted targets capable of supplemental pairing were identified as those with TargetScan7 supplementary pairing scores
< −0.05 (Agarwal 2015).

**Supplemental Figure S5.**
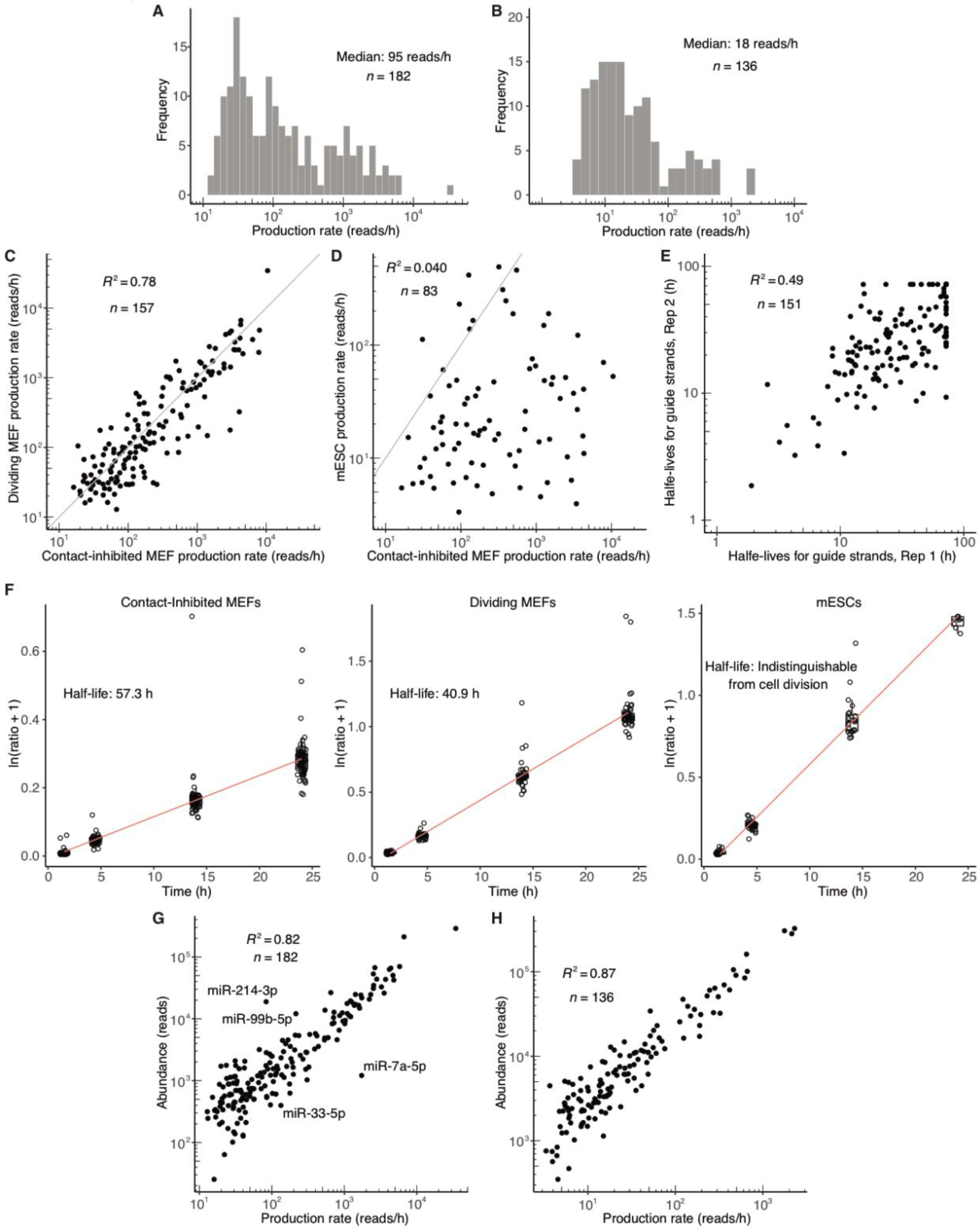
Dynamics of miRNA guide strands in dividing MEFs and mESCs. (A–B) Distribution of production rates for guide strands in dividing MEFs (A) and mESCs (B). (C–D) Relationship between guide-strand production rates in either dividing MEFs (C) or mESCs (D) and guide-strand production rates in contact inhibited MEFs. (E) Comparison of guide strand half-lives for dividing MEF replicates. (F) AGO2 half-lives in contact-inhibited MEFs, dividing MEFs, and mESCs. Shown are plots of the natural-log-transformed ratios of the heavy to light isotopes for all AGO2 peptides identified by mass spectrometry following SILAC pulse-labeling with heavy media in each cell type. These data were fit to a linear function (red) to extract AGO2 half-lives. (G–H) Relationship between guide-strand production rate and miRNA abundance in dividing MEFs (G) and mESCs (H).

**Supplemental Figure S6.**
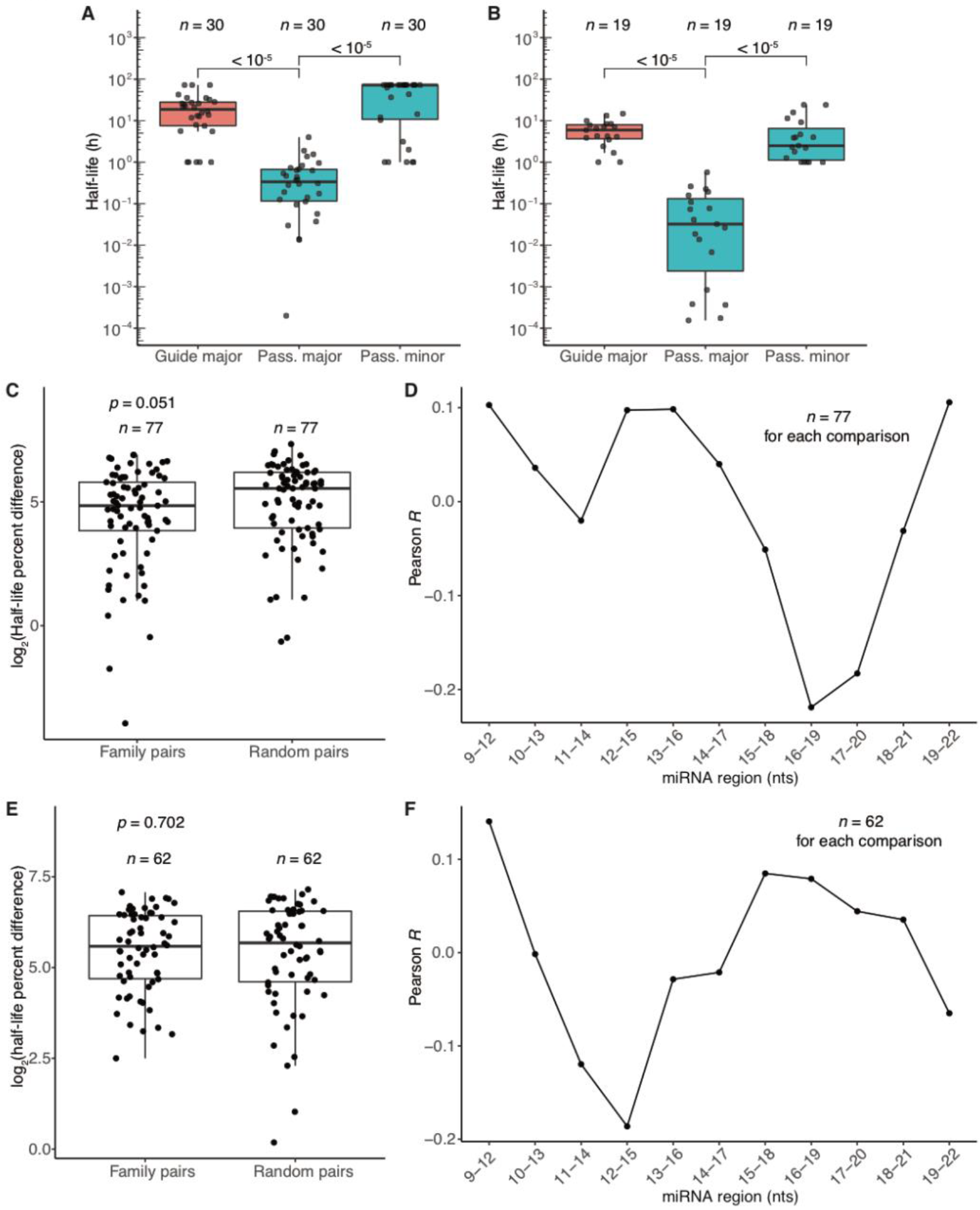
Passenger-strand dynamics and determinants of half-life in dividing MEFs and mESCs. (A–B) Half-life distributions for guide and passenger (pass.) strands of dividing MEFs (A) and mESCs (B), determined by fitting to the biexponential model. Major and minor half-lives are as in Fig. 3B. Significance was determined using a Mann–Whitney test. (C) Differences in half-lives for pairs of family members or randomly paired guides in dividing MEFs. Otherwise, as in Fig. 4A. (D) Relationship between half-life difference and Hamming distance for 4-nt segments of family-member pairs in dividing MEFs. Otherwise, as in Supplemental Fig. S4B. (E) As in (C) but analyzing data from mESCs. (F) As (D) but analyzing data from mESCs.

**Supplemental Figure S7.**
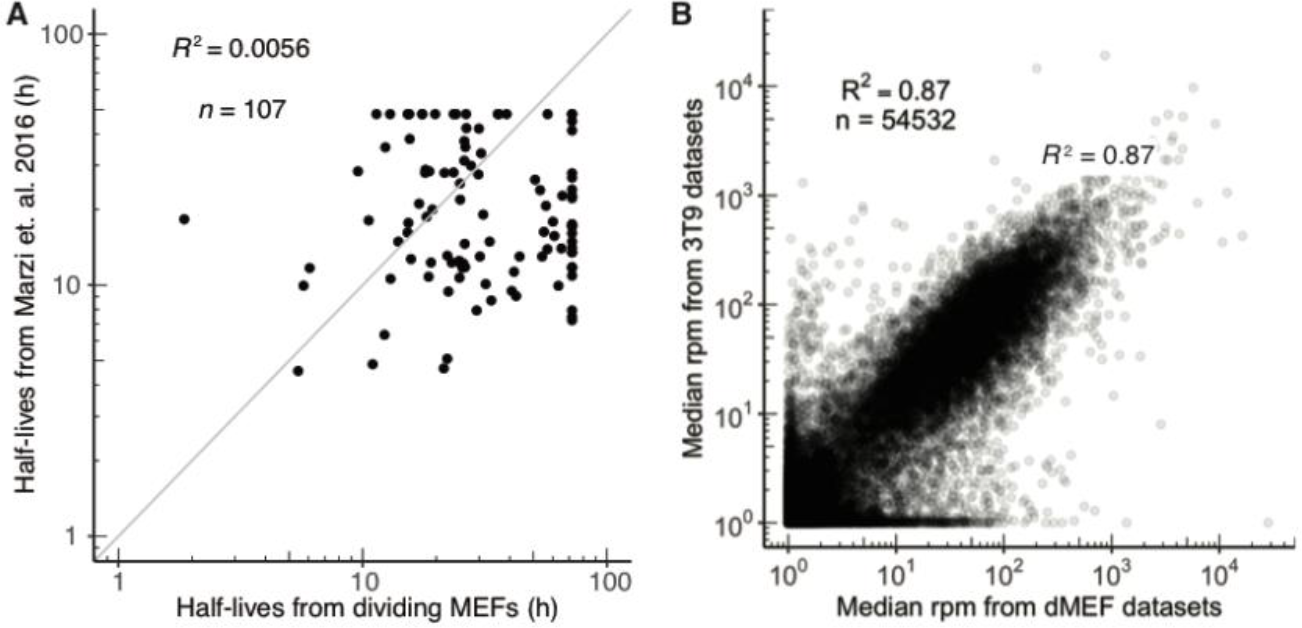
Comparison of published results from dividing 3T9 mouse fibroblasts with our results from dividing MEFs. (A) Correspondence between guide-strand half-lives published for 3T9 cells (Marzi, 2016) and those obtained for dividing MEFs. (B) Correspondence between mRNA levels in dividing 3T9 cells (Ghini, 2018; GEO accession number GSE104650) and mRNA levels in dividing MEFs. Plotted are median RPM values from two and three RNA-seq replicates from 3T9 cells and dividing MEFs, respectively.

**Supplemental Figure S8.**
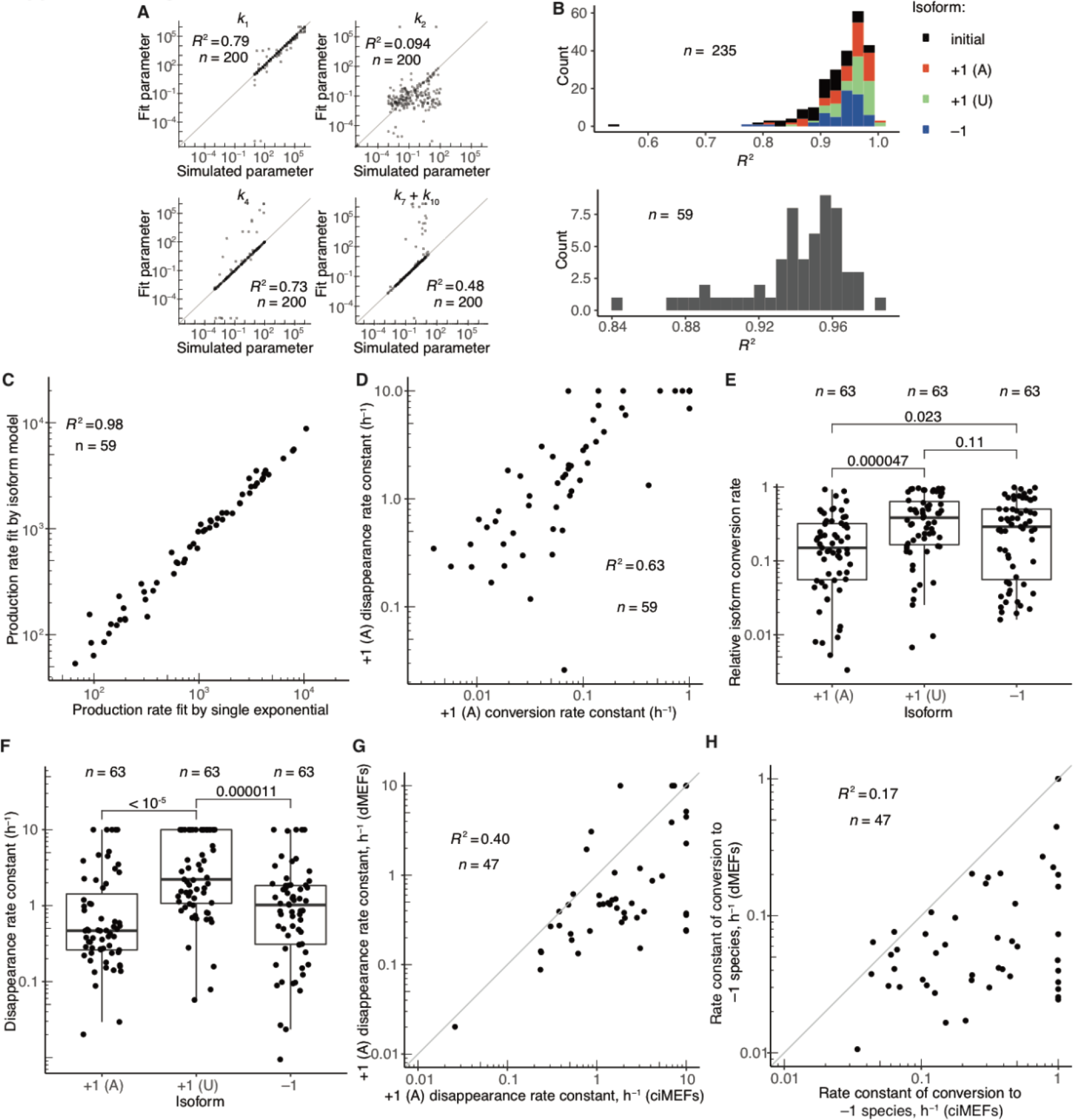
Isoform dynamics in contact-inhibited and dividing MEFs. (A) Correspondence between parameter values fit to simulated data and those initially used to generate the simulated data. A wide array of parameter values was applied to the model described in Fig. 6B to generate the simulated data. Plotted are values fit to these data as a function of the initial simulated values for the rate constant of initial isoform production (*k*_1_), the rate constant of initial isoform degradation (*k*_2_), the rate constant for conversion of the initial isoform to the +1 (U) isoform (*k*_4_), as well as the rate constant for disappearance of the +1 (U) isoform (*k*_7_ + *k*_10_). Although not shown, the correlations for rate constants for the conversion and disappearance of the other isoforms were similar to those observed for +1 (U). (B) Stacked histograms of the *R*^2^ values for fits to each individual isoform (top) as well as all isoforms together (bottom). (C) Relationship between miRNA production rates in contact-inhibited MEFs, comparing the production rate of the initial-isoform fit to the isoform model (Fig. 6B) and the production rate of the guide RNA using the single-exponential fit (Fig. 1B). (D) Relationship between the rate constant of disappearance of the +1 (A) isoform and the rate constant of conversion to the +1 (A) isoform. Because simulations indicated that conversion rate constants >1 h^−1^ and disappearance rate constants >10 h^−1^ could not be accurately fit, conversion and disappearance rate constants were capped at these values, respectively. (E) Relative rates of conversion to the +1 (A), +1 (U), or –1 isoforms in dividing MEFs. Otherwise, as in Fig. 6E. (F) Rate constants of disappearance (*k*_6_ + *k*_9_, *k*_7_ + *k*_10_, or *k*_8_ + *k*_11_) for the +1 (A), +1 (U), or −1 isoform, respectively, in dividing MEFs. Otherwise, as in Fig. 6F. (G) Correspondence between rate constants of disappearance of +1 (A) (*k*_6_ + *k*_9_) in dividing MEFs (dMEFs) and contact-inhibited MEFs (ciMEFs). (H) Correspondence between rate constants of conversion to the −1 isoform (*k*_5_) in dividing MEFs (dMEFs) and contact-inhibited MEFs (ciMEFs).

